# Generative Drug Design in a Loop with dtSFM

**DOI:** 10.64898/2026.06.10.731501

**Authors:** Sai T. Reddy

## Abstract

Directed evolution consisting of iterative rounds of diversification, selection, and counter-selection, underlies modern protein and antibody engineering, yet small-molecule drug design still advances largely through high-throughput screening and medicinal-chemistry intuition. Transformer softmax attention is mathematically identical to the Boltzmann distribution that governs molecular binding at thermal equilibrium^1^, an isomorphism that prescribes a sequence-native Specificity Foundation Model (SFM)^2^. This framework was recently applied across seven molecular recognition domains^3,4^ and scaled into the drug–target SFM (dtSFM), the first to pair a full-scale encoder with a generative decoder^5^. Whether such a model can be driven, iteratively and under selection, to optimize leads rather than sample them once has not been shown. Here we present GenLoop, a closed generative drug design loop that turns single-pass generation into directed evolution of chemistry. dtSFM generates target-conditioned molecules and reranks them by their thermodynamic compatibility score. An orthogonal structural verifier, AlphaFold 3, is used that shares no architecture or training data with dtSFM. Cheminformatics filters enforce developability, and generative evolution is performed on the structurally verified candidates, selecting for predicted binders and counter-selecting against off-target chemistry. Applied across twelve drug targets spanning pharmacologically distinct mechanism classes, GenLoop produced AlphaFold 3-verified designs that reached the structural confidence of the approved drug for five of the twelve targets, with the best designs at interface iPTM 0.93–0.98 and PAE 0.8–2.0 Å, as well as resolving paralog selectivity across nine targets. Two full disease campaigns followed. For the cystic-fibrosis transmembrane conductance regulator, GenLoop designed nine developability-filtered and structurally novel lead candidates (iPTM up to 0.93, interface PAE 2.3 Å) targeting all three orthogonal sites of the approved drug Trikafta. For the GLP-1 receptor family, dtSFM engineered tunable single-, dual-, and triple-receptor incretin designs, yielding 23 central-pocket candidates that are structurally novel at median iPTM 0.89 and interface PAE 1.95 Å. GenLoop with dtSFM brings directed evolution to small molecules through computational-thermodynamic selection; wet-lab validation is the immediate next step.

## INTRODUCTION

Drug discovery begins with the identification of chemical matter that binds a protein target with the required potency, and advances by optimising that matter into a clinical candidate. The standard route runs from high-throughput screening of large compound libraries^6^ or fragment-based discovery^7^ through hit-to-lead and lead optimisation, in which structure-activity relationships, ligand efficiency^8^, and the multi-parameter constraints of absorption, distribution, metabolism, excretion, and toxicity^9^ are balanced over successive rounds of synthesis and experimental assays. Selectivity is decisive throughout: a candidate must engage its intended target while avoiding the closely related proteins (paralogs) and the standard off-target panels, including kinase families, the hERG channel, and the cytochrome-P450 enzymes, whose inadvertent engagement drives attrition and clinical toxicity^10,11^. The accessible space of drug-like molecules is vast, estimated near 10^60^ ^12^, and experimental discovery is bounded by the membership of the libraries that are actually synthesized.

Computational drug design has long approached binding through three-dimensional structure. Molecular docking and structure-based virtual screening rank candidate poses against a target pocket^13^, and physics-based methods estimate binding free energies directly through molecular dynamics^14^ and free-energy perturbation^15^. Generative methods extend this structural paradigm: structure-first models place atoms within a defined pocket, including the diffusion and autoregressive generators DiffSBDD^16^, Pocket2Mol^17^, and TargetDiff^18^, while sequence-native models generate the molecular string itself, in the lineage of the SMILES language models MolGPT^19^ and Chemformer^20^. The structural verification of a designed candidate has in turn been transformed by protein-ligand cofolding: AlphaFold 3 (AF3) ^21^ and the open-source Boltz-1 and Boltz-2 models^22,23^ predict the geometry of a drug-target complex and report calibrated interface-confidence metrics, providing a structural readout independent of the model that generated the candidate.

We take a different approach, grounded in the mathematical isomorphism: the softmax attention of transformers^24^ is the Boltzmann distribution governing molecular binding at thermal equilibrium, which we term the convergence equation and from which the architecture of a Specificity Foundation Model (SFM) is prescribed along with the framework for Molecular Recognition Computing (MRC)^1–3^. An SFM does not approximate an unknown function; it estimates the parameters of the convergence equation that governs binding, performing computational thermodynamics by ranking binding as a free-energy-like compatibility computed directly from the sequences of the two partners, without ever constructing a three-dimensional coordinate. The drug-target SFM (dtSFM) realized the SFM for small molecule drug-target protein domain and reported the first trained instance of the cross-attentive SFM decoder, a target-conditioned generator that samples chemistry from the thermodynamic energy landscape the encoder defines^5^. Applied once to sixteen targets, the dtSFM decoder matched the structural confidence of the approved drug on a majority of its designs, yet the per-target yield was heterogeneous, and a single generative pass addressed neither selectivity against related proteins, nor depth across multiple pockets on one target, nor the tunable polypharmacology that distinguishes modern therapeutics.

Here we present GenLoop: a closed, iterative generative drug design and evolution loop built on dtSFM, that converts single-pass generation into multi-parameter-optimised lead cohorts. Within the loop, dtSFM generates target-conditioned candidate molecules and reranks them by their thermodynamic compatibility score; an orthogonal protein-ligand cofolding model (e.g., AF3) provides the structural verification step and is independent of the dtSFM scoring axis; a stack of cheminformatics filters enforces synthesizability, drug-likeness, and scaffold novelty; and a generative evolution refinement stage optimises the surviving cohort, either via bioisostere mutagenesis libraries built from a verified parent scaffold, the chemical analog of single-site and combinatorial mutagenesis, and by low-rank (LoRA) fine-tuning of the decoder toward the validated pocket. We apply GenLoop to design drug candidates across four settings: structure-verified breadth across twelve mechanism classes, paralog-resolved selectivity, multi-site depth on the three orthogonal sites of the cystic fibrosis transmembrane conductance regulator CFTR, and tunable single-, dual-, and triple-receptor polypharmacology on the GLP1-family receptors. In concept GenLoop is directed evolution applied to small molecules, a generated library that is structurally selected and counter-selected and refined over successive rounds, operating within dtSFM that computes molecular recognition as thermodynamic selection.

## RESULTS

### GenLoop is directed evolution of small molecules with dtSFM

Directed evolution is a well-established method in protein engineering for improving the binding and functional properties of antibodies, enzymes, and other proteins^25^. It proceeds through iterative rounds in which sequence variants are generated by single-site or combinatorial mutagenesis, screened by display platforms for on-target binding or function, counter-selected against off-target reactivity, and advanced through lead optimisation steps that address multiple parameters in parallel, including binding, specificity, and developability. We have previously established this approach in two molecular recognition systems. In antibody engineering, mutagenesis screening of a therapeutic antibody by mammalian display against its antigen was deep-sequenced to train a neural network that extrapolated to approximately 7 × 10⁷ in silico variants, jointly filtered for binding and multiple developability properties^26^. In T cell receptor engineering, combinatorial mutagenesis libraries of approximately 2.6 × 10⁵ variants were screened for antigen-induced functional activation, with counter-selection against off-target reactivity^27^. Two principles transferred from both systems: the model’s scoring axis must be paired with a selector that measures function orthogonally to it, rather than predicted affinity per se; and multi-parameter optimisation is the output of the loop, not of any single round.

GenLoop is the small-molecule analog of these directed-evolution platforms, built on dtSFM^5^, and comprises four steps (**Fig. 1**). The first step replaces experimental library generation with autoregressive SMILES^28^ sampling from the target-conditioned chemical decoder of dtSFM in the lineage of MolGPT-style chemical language models^19^; the dtSFM decoder is sampled at temperature T = 0.8, EOS-terminated, with a 120-token cap, producing n = 512 raw candidates per target as a default budget. The second step replaces a pre-sort step with the dtSFM encoder rerank based on the drug–target cosine similarity, which sorts the raw pool and forwards the top 20 % to structural verification. The third step replaces the orthogonal functional selector (mammalian or yeast display, flow cytometry, and next-generation sequencing) with AF3 cofolding of each candidate against the intended target; each cofold returns an interface predicted TM score (iPTM) and an interface predicted aligned error (PAE), and candidates are classified against decoy-calibrated joint thresholds: STRONG, MODERATE, and WEAK by joint interface iPTM and PAE (thresholds calibrated below). The fourth step is generative evolution refinement of each target’s AF3-verified positive cohort, in two complementary modes detailed below: a generative bioisostere mutagenesis library, and LoRA-based fine-tuning of the dtSFM decoder^29^ paired with an unlikelihood loss^30^ on off-site and paralog regressors that takes the role of negative panning or off-target counter-selection^27^. The multi-parameter developability filter that follows each generation is computed in RDKit^31^ and combines SMARTS-encoded reactive-group alerts^32^ drawn from the Brenk panic-alert library^33^, pan-assay-interference (PAINS) flags^34^, the Lipinski Rule-of-Five (Ro5) developability gate^9^, the quantitative estimate of drug-likeness (QED)^35^, synthetic-accessibility scoring^36^, Bemis–Murcko scaffold extraction^37^, and structural novelty filtering by Morgan extended-connectivity (ECFP4) Tanimoto distance to the clinical anchors of each campaign^38^. The orthogonality of the encoder cosine and the AF3 interface confidence, established in the preceding dtSFM paper^5^ underwrites GenLoop’s use of the encoder cosine and the AF3 interface confidence as orthogonal filters.

**Figure 1.**
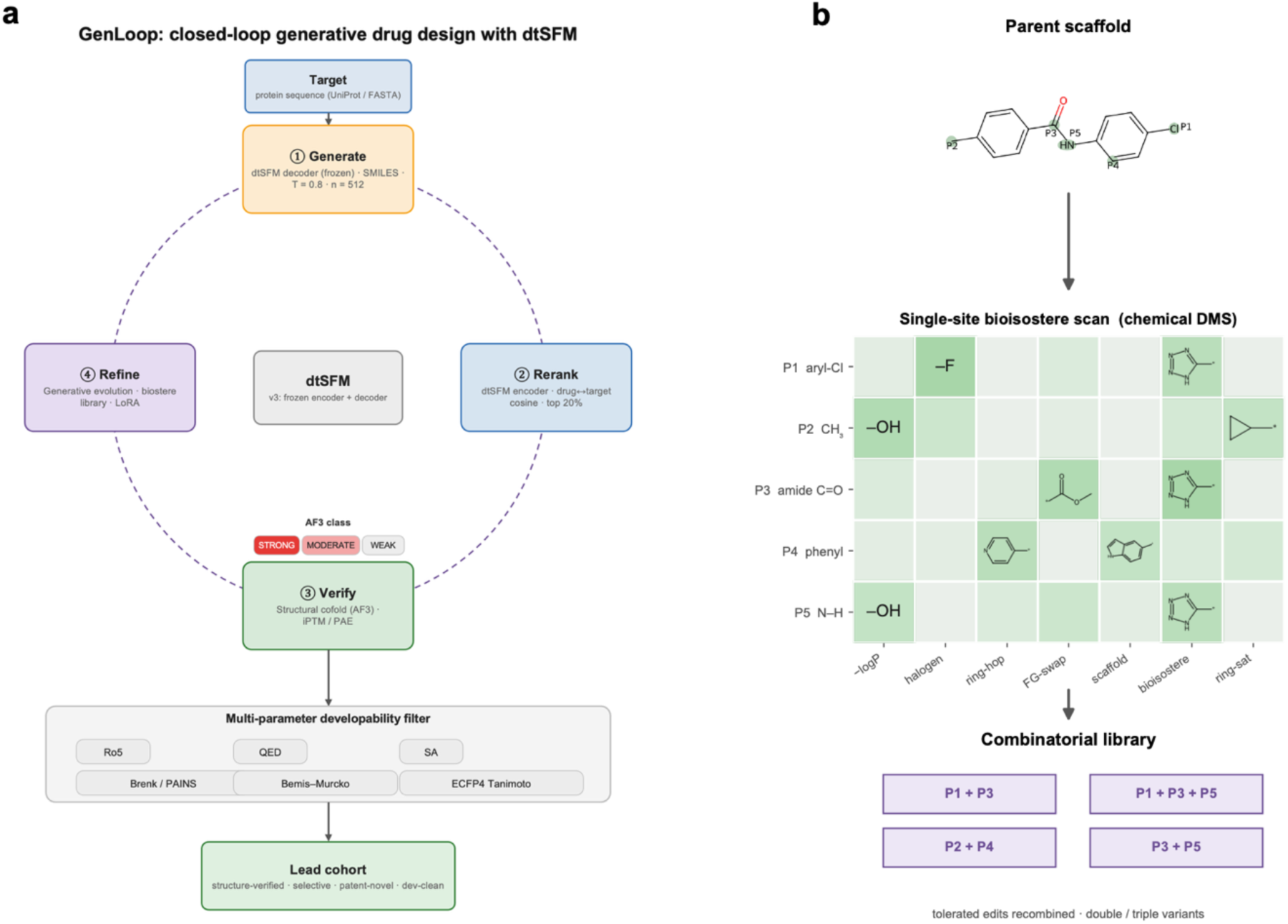
Generative drug design in a loop with dtSFM (GenLoop). **(a)** dtSFM drives a closed, four-step generative loop. **1**. Generate: a target-conditioned chemical decoder (dtSFM decoder v3, 27 M parameters, frozen base) samples SMILES candidates from the target sequence (temperature T = 0.8, n = 512 per target). 2. Rerank: the dtSFM encoder scores each candidate by drug-target cosine and forwards the top 20 %. 3. Verify: AlphaFold 3 (AF3) cofolds the reranked candidates against the target and emits iPTM and interface PAE, classifying each as STRONG / MODERATE / WEAK against decoy-calibrated thresholds. 4. Refine: AF3-verified candidates seed a generative evolution refinement stage — bioisostere mutagenesis and/or LoRA fine-tuning of the decoder on the structurally verified cohort. Verified candidates then pass a multi-parameter developability filter (Ro5, QED, synthetic accessibility, Brenk / PAINS alerts, Bemis–Murcko scaffolds, ECFP4 Tanimoto novelty) resulting in a lead cohort. **(b)** Library design for generative evolution. From an AF3-verified parent scaffold, a curated set of about 80 medicinal-chemistry bioisostere SMARTS transforms is applied, introduced as a single-site scan (chemical analog of saturation deep mutational scanning) or in combination (the analog of combinatorial mutagenesis), stepwise or in parallel. Single-site scanning first identifies the tolerated edits, focusing a subsequent combinatorial library on viable chemistry; library members are scored and selected with the dtSFM encoder and AF3.

GenLoop performs *generative evolution*: it diversifies and selects chemical matter the way directed evolution diversifies and selects proteins, except that the diversity is generated in silico and the selection is computed as thermodynamic compatibility rather than recombinantly expressed and assayed. Generative evolution proceeds through mutagenesis libraries that can be designed in more than one way. Starting from a parent scaffold the loop has already verified against the target, we apply a curated set of about 80 medicinal-chemistry bioisostere transforms encoded as RDKit SMARTS rules (lipophilicity-reducing edits, halogen swaps, ring and scaffold hops, functional-group and bioisosteric replacements). These edits can be introduced one at a time, a single-site scan that is the chemical analog of saturation deep mutational scanning^39^ or several edits at once, the analog of combinatorial mutagenesis^26,27,40^, and the two can be run stepwise or in parallel. The single-site scan is most valuable when downstream screening capacity is limited: by first identifying which single substitutions the target tolerates, it focuses a subsequent combinatorial library on the positions and chemistries the target accepts, so that fewer combinations need to be made and tested. Because GenLoop generates and scores every library member computationally, with the dtSFM encoder and AF3, it can apply whichever design fits the campaign, and at a scale a conventional medicinal-chemistry program cannot reach, which requires each analog to be individually synthesized and assayed through design-make-test cycles^41^. The selected library members re-enter the loop (**Fig. 1B**).

### GenLoop generates AF3-verified candidates across 12 mechanism classes

We applied GenLoop to a panel of 12 drug targets spanning pharmacologically distinct mechanism classes, including kinases ALK, BTK, JAK3, and TYK2, the nuclear receptor THRB, the complement protease CFB, the protein–protein interaction target MEN1 (menin–MLL), the phosphatase SHP2, the inflammasome sensor NLRP3, and three Class B G-protein-coupled receptors (GCGR, GIPR, GLP1R). For each target, the base decoder produced n = 512 raw samples at temperature T = 0.8, of which 53 % (GLP1R) to 76 % (ALK) parsed as valid molecules under RDKit and 51 % to 75 % were uniquely canonicalised (**Fig. S1a**); pooled across all 12 targets this yielded n = 61,766 raw decoder candidates entering the loop. The pooled starting chemistry had a median QED of 0.37 and a median Tanimoto of 0.18, and 43 % of candidates were Ro5 (**Fig. 2a**); per-target Ro5 fraction ranged from approximately 0 % (GCGR) to approximately 80 % (NLRP3) (**Fig. S1b**). Gating the top 20 % of each per-target pool by the dtSFM drug–target cosine preserved median novelty (0.82 → 0.78) while focusing scaffold diversity (Bemis–Murcko: 0.43 → 0.31), confirming that the encoder rerank operates as a relevance triage rather than a novelty-collapsing filter (**Fig. 2b**).

**Figure 2.**
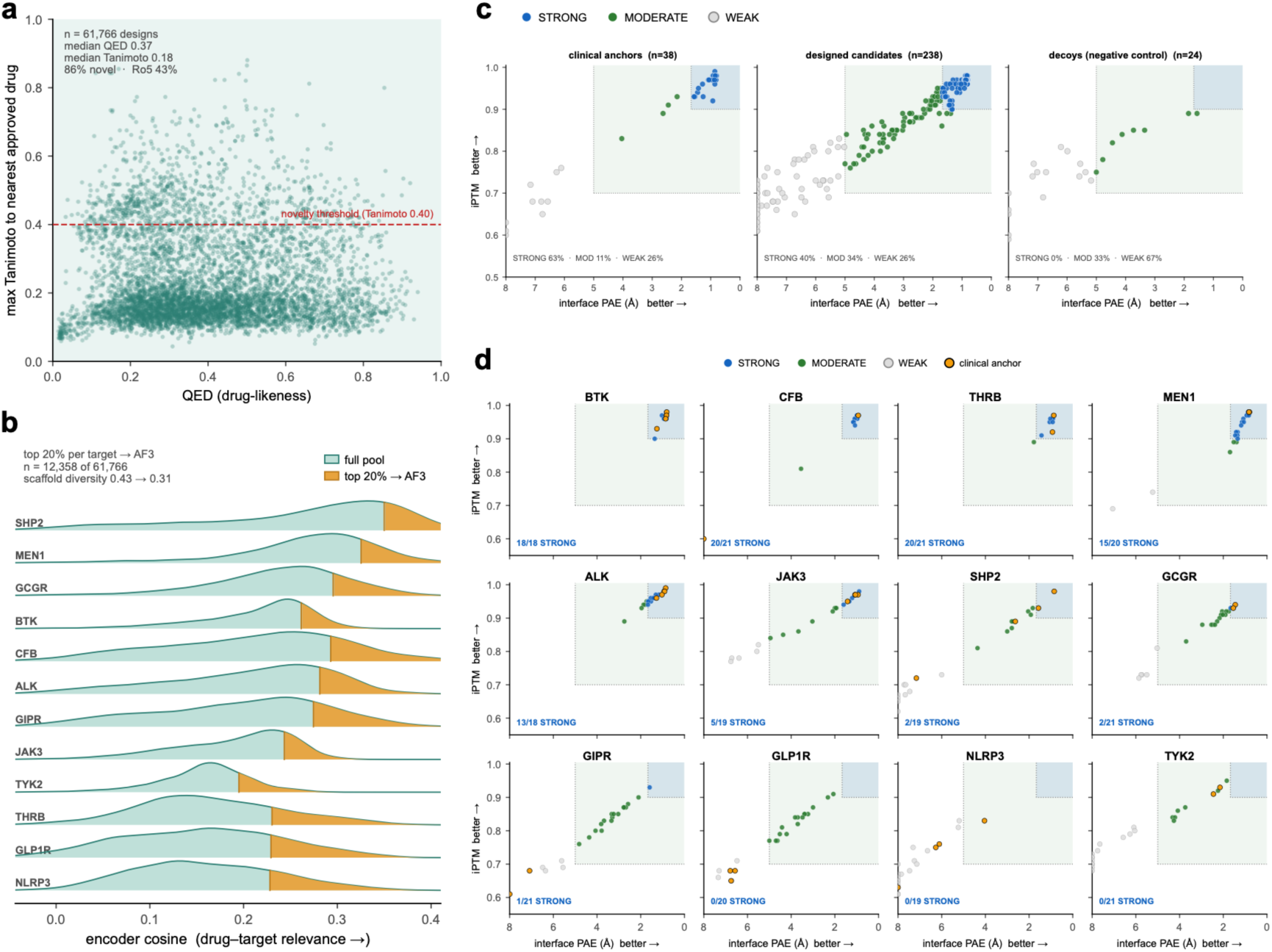
Decoder → encoder → AF3 screening across 12 breadth targets. **(a)** Starting-pool quality. n = 61,766 raw decoder candidates pooled across 12 breadth targets, plotted as drug-likeness QED versus ECFP4 Tanimoto similarity to the nearest approved drug. Median QED = 0.37; median Tanimoto = 0.18, with 86 % of candidates below the 0.40 novelty threshold (dashed line); approved-drug self-similarity (Tanimoto = 1.0). 43 % Lipinski Ro5-compliant. **(b)** Encoder rerank as a per-target relevance gate, shown as a ridgeline of raw dtSFM drug–target cosine across the 12 breadth targets (one density per target, sorted by median cosine). Each target’s pool spans a broad, target-specific cosine range; GenLoop applies one uniform top-20 % gate per target (gold tail), carrying 12,358 of 61,766 candidates forward to AF3. The rerank focuses scaffold diversity only modestly (Bemis–Murcko unique scaffolds per candidate: 0.43 → 0.31. **(c)** AF3 validation of three populations gated identically (n-design = 238; n-anchor = 38 clinical / research-stage references; n-decoy = 24 negative-control molecules). Designed candidates: 40 % STRONG / 34 % MODERATE / 26 % WEAK. Anchors: 63 / 11 / 26 %. Decoys: 0 / 33 / 67 %. **(d)** Per-target screening landscape. All 12 targets shown as gate plots of AF3 iPTM (vertical, better → top) versus reversed interface PAE (horizontal, better → right). Designed candidates coloured by class (STRONG = blue, MODERATE = green, WEAK = light gray); clinical anchors overlaid as gold circles. Per-target STRONG count annotated. Panels sorted by % STRONG.

We calibrated the AF3 confidence gate empirically on a panel of n = 24 negative-control molecules drawn from drug-like SMILES with no documented relationship to any target in the 12-target screen, by cofolding each decoy against each of the 12 targets in AF3. Cohort-wise decoy pass-rates were 33 % at the MODERATE gate (iPTM ≥ 0.70 and interface PAE ≤ 5 Å) and 0 % at the STRONG gate (iPTM ≥ 0.90 and interface PAE ≤ 1.67 Å) (**Fig. S2**). The STRONG gate is therefore the empirically calibrated specific-binding threshold of the platform, and the MODERATE gate is reported transparently as ambiguous (within the AF3 noise floor) throughout this paper. Evaluated on the same gate, three populations cofolded against the 12-target panel separated cleanly (**Fig. 2c**): designed candidates (n = 238 top-rerank candidates carried into structural verification) split 40 % / 34 % / 26 % into STRONG / MODERATE / WEAK; clinical and research-stage anchors (n = 38) split 63 % / 11 % / 26 %; and the 24 decoys split 0 % / 33 % / 67 %, respectively. Designed candidates reached STRONG fractions within the same order as published clinical anchors and an order of magnitude above the empirical decoy noise floor.

The per-target screening landscape resolved each of the 12 targets as an AF3 gate plot of iPTM versus reversed interface PAE (**Fig. 2d**). Stratifying the 12 targets by STRONG fraction (**Table S1**) revealed three tiers: five targets reached ≥ 70 % STRONG (BTK 100 %, THRB 95 %, CFB 95 %, MEN1 75 %, ALK 72 %); four targets reached partial yield of 5–30 % STRONG (JAK3 26 %, SHP2 11 %, GCGR 10 %, GIPR 5 %); and three targets returned 0 % STRONG (TYK2, NLRP3, GLP1R). The three zero-yield targets are not failures of the platform but of its single-pass screen, and the generative evolution refinement step of GenLoop, developed for GLP1R in the polypharmacology subsection demonstrates recovery.

### Structural characterization of dtSFM-generated variants

We focused structural inspection on all five STRONG-tier targets from the breadth screen (BTK, CFB, THRB, ALK, MEN1). For each target, the AF3-predicted complex of the top-9 designed candidates was cealigned against the AF3-predicted complex of the clinical anchor on the target’s binding pocket (**Fig. 3**). For THRB / resmetirom and MEN1 / revumenib, a nuclear receptor and a protein-protein interaction inhibitor, the designed candidates occupied the same pocket as the clinical anchor with matched AF3 confidence (design iPTM 0.93 – 0.98, interface PAE 0.8 – 2.0 Å) while remaining chemically distinct (Tanimoto 0.16 – 0.25 to the anchor). ALK occupied the lorlatinib pocket with comparable confidence and novel scaffolds, but most of its top designs carry reactive alkyl-halide termini (ALK 11 % chemistry-clean; **Table S1**). A single-pair zoom for THRB, ALK, and MEN1 together with the cleanest JAK3 design forms a four-cell representative gallery (**Fig. S3a**), with the corresponding 2D chemistry pairs (**Fig. S3b**) and full per-candidate detail (**Table S2**). The two remaining columns report decoder pathologies (**Fig. 3**): all 18 BTK STRONG designs carry a reactive vinyl-halide warhead learned from the covalent anchors and are readily synthesizable (synthetic-accessibility 2.7–4.2) but flagged for reactivity by the Brenk structural-alert filter^33^, and the top-9 CFB STRONG designs are iptacopan-like (Tanimoto 0.57–0.87, decoder memorisation rather than scaffold novelty).

**Figure 3.**
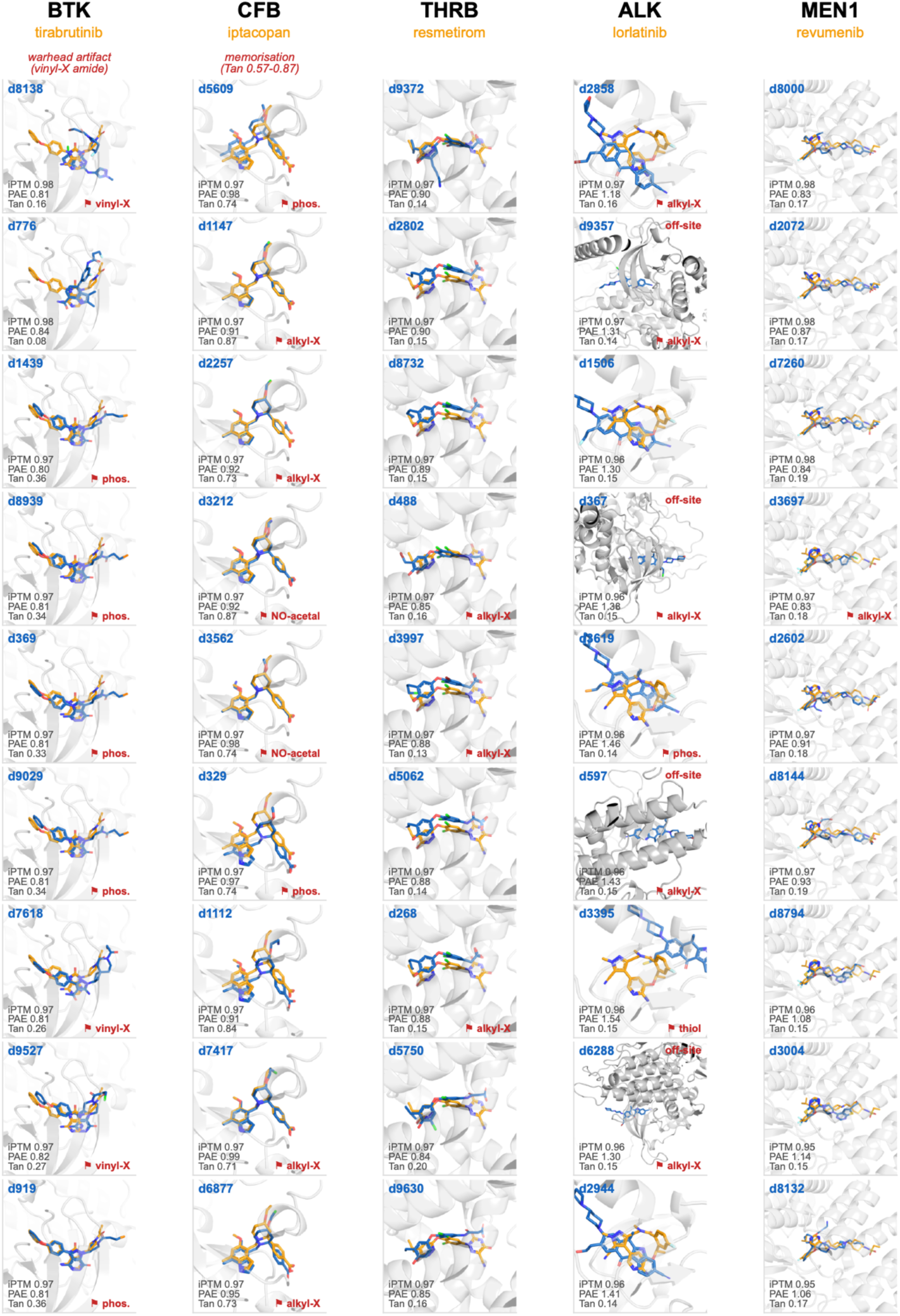
Structural wall of AF3-predicted overlays across five STRONG-tier breadth targets. Per-target AF3-predicted complexes of the top-9 designed candidates per target (rows) cealigned against the clinical anchor’s AF3-predicted complex on the target’s binding pocket. Columns are the five STRONG-tier targets ordered by per-target % STRONG: **BTK** (anchortirabrutinib), **CFB** (iptacopan), **THRB** (resmetirom), **ALK** (lorlatinib), **MEN1** (revumenib). Each cell shows the designed candidate (blue sticks) cealigned to the clinical anchor (gold sticks) on the target pocket (semi-transparent gray cartoon), sharing a single tight per-target pocket camera so candidates are directly comparable down a column. Cell labels: designed candidate identifier (dNNNN, top left, blue); AF3 iPTM and interface PAE (Å), and Tanimoto similarity to the anchor (bottom left). **BTK** carries a warhead-artifact tag (all 18 BTK STRONG designs share a Michael-acceptor vinyl-X amide motif with a non-synthesizable terminal atom, learned from the two covalent clinical anchors); **CFB** carries an iptacopan-memorisation tag (Tanimoto 0.57–0.87 to the anchor across all top-9 designs). Cells whose ligand sits > 20 Å from the anchor pocket post-cealign are labelled “off-site” (top right, red).

Across the five STRONG-tier targets, a tightened reactive-group filter (primary phosphines, alkyl and vinyl halides, halo-ethers, N,O-acetals, free thiols; carbon-bound fluorine, being metabolically stable, is not flagged) leaves 29 of 86 STRONG designs (34 %) free of structural-alert flags (**Table S1**). The clean fraction tracks how faithfully the decoder reproduced the anchor’s stable chemistry — high where it did (THRB 62 %, MEN1 55 %), low where it collapsed onto a reactive terminal motif (BTK 11 %, CFB 5 %, ALK 11 %). AF3 structural confidence is a readout of pocket geometry, not chemical stability: a reactive primary phosphine cofolds as confidently as a clean scaffold. The STRONG gate is therefore a structural filter, completed downstream by the cheminformatics developability gates and, where a motif recurs, by generative evolution refinement. This is the intended output of a breadth screen — a chemically diverse pool of target-matched scaffolds carrying precisely the liabilities that a lead-optimisation phase exists to resolve, as in any hit-to-lead campaign.

### Structure-based paralog selectivity profiling of dtSFM-generated variants

Before reporting per-target selectivity outcomes, we first verified that the AF3 paralog cofold readout itself discriminates intended from off-target binding. Across the n = 205 paralog cofolds carrying both scores, AF3 was on average 1.22 Å more confident on the intended target than on the paralog, while the iPTM difference was essentially zero (mean Δ-PAE = +1.22 Å, mean Δ-iPTM = −0.08, paralog − intended; **Fig. S4a**). Interface PAE therefore discriminates selectivity that iPTM does not and is the axis on which the per-target tiering below is defined.

For each target we assembled a paralog panel from the closest sequence and structural homologues within its gene family, the proteins against which clinical selectivity is most stringently required and most often lost, exemplified by the human kinome for the kinase targets^42^ and the incretin and Class B GPCR families for the receptor targets. We profiled paralog selectivity on the 9 of 12 breadth targets carrying such panels (BTK, CFB, THRB, ALK, MEN1, GCGR, GIPR, GLP1R, TYK2; 3 to 6 paralogs each, n = 41 paralogs in total); we chose not to profile the three targets without curated paralog panels (JAK3, NLRP3, SHP2; **Fig. S4b**). For each of the 9 targets, all designed candidates and the clinical anchor were cofolded in AF3 against the intended target and against each member of the paralog panel, the resulting cofolds were classified at the decoy-calibrated joint thresholds defined in the breadth subsection (STRONG / MODERATE / WEAK). We assigned each target an operational selectivity tier based on the fraction of its paralog cofolds reaching STRONG: CLEAN (< 25 % of paralog cofolds STRONG), PARTIAL (25–75 % STRONG), and FAMILY-SHARED (> 75 % STRONG). Of the 9 targets, 5 were CLEAN (GCGR, GIPR, GLP1R, MEN1, TYK2), 3 were PARTIAL (ALK, CFB, THRB), and 1 was FAMILY-SHARED (BTK) (**Fig. 4a** and **Table S3**).

**Figure 4.**
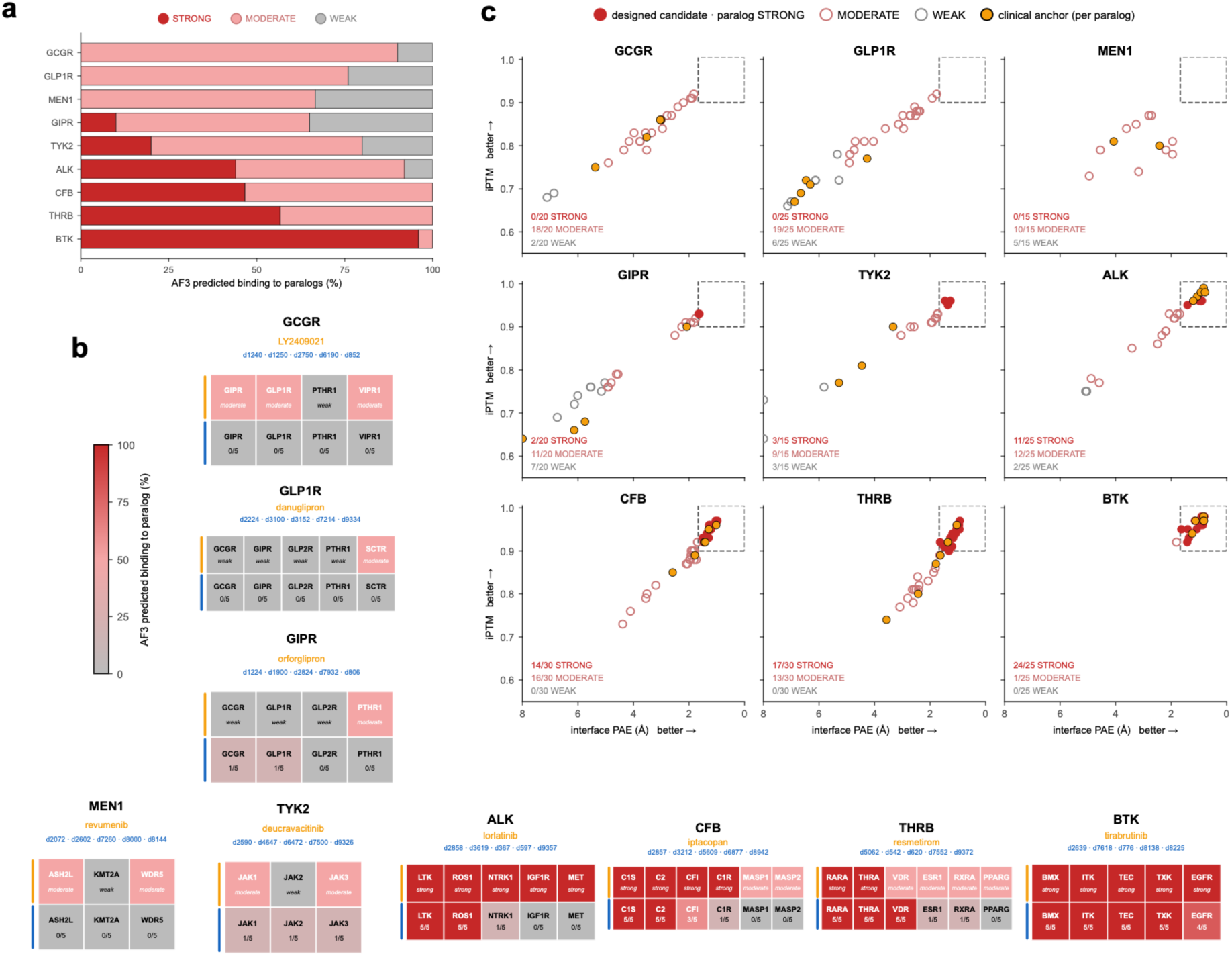
Paralog selectivity across 9 breadth targets. Selectivity tiers, operational: **(a)** Per-target paralog-binding spectrum. Each row stacks the fraction of that target’s paralog cofolds by AF3 class (STRONG = red, MODERATE = salmon, WEAK = gray); rows sorted top → bottom from most-selective to family-shared. **(b)** Cross-reactivity matrix. Each card = one target; card header lists the target name (black), the clinical anchor (gold), and the dNNNN identifiers of the 5 dtSFM-designed candidates whose cofolds populate the card (blue). Below the header, each card is a two-row mini-heatmap, one column per paralog: top row = clinical anchor’s AF3 class on each paralog; bottom row = designed candidates’ fraction-STRONG (n-STRONG / n-total). Both rows share a gray → salmon → red colour scale. **(c)** Per-target paralog gate plots (iPTM vs reversed PAE), one panel per target; designed-candidate paralog cofolds coloured by class (STRONG = filled red, MODERATE = open salmon, WEAK = open gray); clinical-anchor paralog cofolds overlaid as gold filled circles.

A central finding emerged on the FAMILY-SHARED and PARTIAL targets (**Fig. 4b**): on the conserved-fold targets, the clinical anchor itself cross-reacted on the same paralogs the designs hit: lorlatinib was STRONG on 5 of 5 ALK-family paralogs, tirabrutinib on 5 of 5 BTK-family paralogs, iptacopan on 4 of 6 complement-protease paralogs, and resmetirom on 2 of 6 THRB nuclear-receptor paralogs. The FAMILY-SHARED outcome is therefore a property of the conserved fold rather than a failure of the design method. The reciprocal observation also held: the incretin-class clinical anchors (orforglipron, danuglipron, LY2409021) and the JAK-selective deucravacitinib reached zero STRONG paralog cross-reactivity, mirroring their clinical selectivity profiles, and the dtSFM-generated variants at these targets were CLEAN alongside the anchors (**Fig. 4c**).

### Generative design and evolution of lead candidates for cystic fibrosis therapy

Cystic fibrosis is an autosomal recessive genetic disease caused by mutations in the gene encoding the cystic fibrosis transmembrane conductance regulator (CFTR), a chloride channel at the apical membrane of epithelial cells. Loss of CFTR function impairs chloride and fluid transport, producing thick airway mucus, chronic pulmonary infection, progressive respiratory decline, and pancreatic insufficiency. The most effective approved therapy is Trikafta, the triple combination of elexacaftor, tezacaftor, and ivacaftor^43^, which acts on CFTR through three orthogonal mechanisms at three distinct sites on the channel: ivacaftor is a potentiator that increases the open probability of channels already at the membrane (site A), while tezacaftor (a Type II corrector, site B) and elexacaftor (a Type III corrector, site C) improve the folding and trafficking of mutant CFTR to the cell surface.

We ran GenLoop against CFTR as a single target and let the binding sites emerge. In the first round the decoder generated candidates, the dtSFM encoder reranked them by cosine similarity to CFTR, and the top-ranked candidates were cofolded against CFTR in AF3 and assigned to a pocket by gemmi canonical-frame alignment^44^. This first round recovered all three Trikafta pockets without any site being imposed during generation, and it identified a shortlist of 34 verified parent scaffolds spread across them. The clinical anchors provide a same-fold reference for these AF3 scores: ivacaftor (iPTM 0.73, interface PAE 6.94 Å) and elexacaftor (iPTM 0.73, interface PAE 7.93 Å) cofold only within the WEAK band, while tezacaftor reaches the STRONG/MODERATE boundary (iPTM 0.92, interface PAE 2.40 Å) (**Fig. 5a**). Because two of the three approved drugs score WEAK on their own target, we evaluated designs by parity with these clinical references rather than against an absolute STRONG gate.

**Figure 5.**
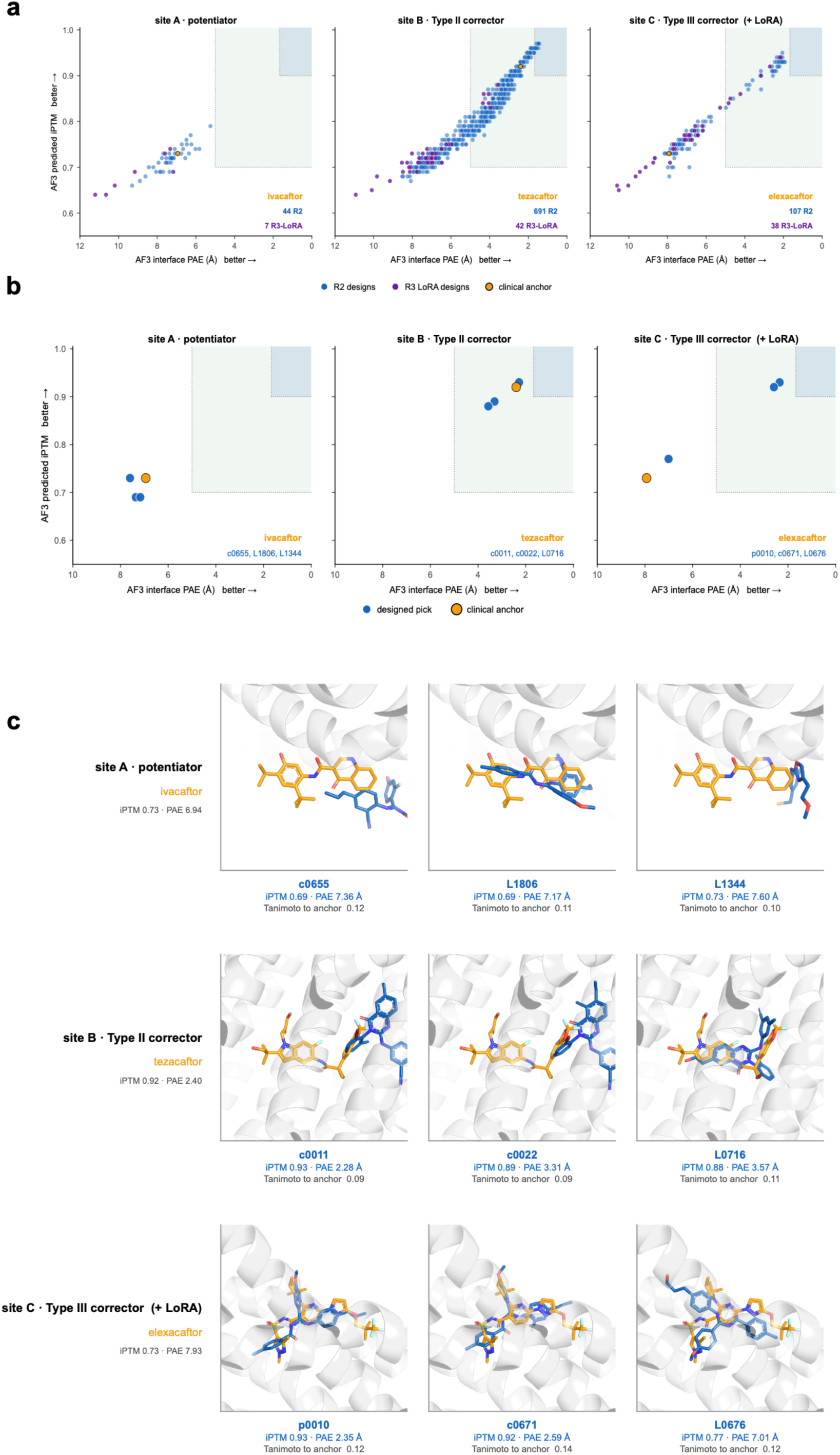
Depth on CFTR: three orthogonal Trikafta sites. CFTR is treated clinically with three orthogonal small molecules — **ivacaftor** (potentiator, site A), **tezacaftor** (Type II corrector, site B), **elexacaftor** (Type III corrector, site C). GenLoop is run separately for each site, with a site-C LoRA-refinement round (**Fig. S6**). **(a)** Full AF3-verified pool per site, plotted as gate plots of iPTM (vertical, better → top) vs reversed interface PAE in Å (horizontal, better → right). Shaded bands mark the STRONG and MODERATE gates. Base-decoder R2 designs are blue circles; LoRA-refined R3 designs are purple circles; the clinical anchor for each site is a gold circle. n = 929 cofolds (site A 51, site B 733, site C 145). **(b)** AF3 verification of the 9 lead candidates (3 per site), same axes as (a) zoomed to the 9 picks plus their anchors. **(c)** Structural overlay gallery of AF3-predicted complex of each designed candidate (blue sticks) cealigned to its site’s clinical anchor (gold sticks) on CFTR (semi-transparent gray cartoon). Each cell labels the design’s c/p/L identifier, its AF3 iPTM and PAE (Å), and its Tanimoto to the anchor; row labels (left) name the site and the anchor’s own AF3 iPTM and PAE.

These 34 parents seeded a round of generative evolution. Applying the bioisostere transforms in combination to the verified parents produced an 845-member combinatorial library that re-entered the loop, with each member rescored by the dtSFM encoder cosine to CFTR and cofolded against CFTR in AF3.

This first combinatorial round under-sampled the elexacaftor pocket: only 7.5 % of cofolded candidates achieved a strict gemmi canonical-frame site match to site C, the deepest and most structurally distinct of the three pockets. To bias generation toward this site, we applied a round of generative evolution refinement using a LoRA adapter^29^ (688 K trainable parameters, 2.5 % of the 27 M-parameter dtSFM v3 decoder) attached to the decoder and trained on 344 site-resolved molecules: 61 site-C-verified positives (combinatorial-round candidates that AF3 placed at the elexacaftor pocket by gemmi assignment) and 283 negatives (candidates placed at sites A or B), under an unlikelihood loss^30^ that pushes the decoder toward the site-C positives and away from the off-site negatives (**Fig. S6a, b**). The refinement raised the fraction of the cofolded pool with strict site-C match from 7.5 % (combinatorial round, n = 842) to 36.0 % (LoRA-refined round, n = 89), a 4.8-fold enrichment, while preserving AF3 confidence on the elexacaftor pocket (median iPTM 0.78 to 0.77, median interface PAE 6.61 to 6.87 Å; **Fig. S6c, d**). The site-C objective shifts the decoder’s output distribution rather than restricting it: the refined samples still populate all three pockets (**Fig. 5a**), and several of the synthesis-clean leads carried forward at the potentiator and Type-II corrector sites come from this round. The refinement is therefore a site-occupancy operation that concentrates generation on the hardest pocket without changing the AF3 confidence achievable there. Pooling the combinatorial and refinement rounds gave 929 verified candidates assigned across the three pockets (by nearest-anchor assignment, 51 at site A, 733 at site B, and 145 at site C; Fig. 5a), the pool from which leads were selected.

We selected three lead candidates per site (nine in total) from this AF3-verified pool after passing each through the full cheminformatics developability stack (Brenk SMARTS reactive-group alerts, PAINS flags, Ro5, QED, synthetic-accessibility scoring, and Bemis-Murcko scaffold novelty), shown alongside their anchors with full per-candidate detail (**Fig. 5b**; **Table 4**). The cohort draws on both rounds, with combinatorial and LoRA-refined designs among the leads at all three pockets. These candidates are strong leads for experimental follow-up for three reasons. First, they reach the AF3 confidence of the corresponding clinical anchor at each site: the site B leads (iPTM 0.88-0.93) match tezacaftor, and the site C leads p0010 (iPTM 0.93, interface PAE 2.35 Å) and c0671 (iPTM 0.92, interface PAE 2.59 Å) exceed the WEAK elexacaftor anchor on both metrics. Second, they are chemically distinct from the anchors, all nine sitting at Tanimoto 0.09-0.14 to the corresponding clinical molecule. Third, they cleared the synthesizability and drug-likeness gates that define a tractable starting point for medicinal chemistry, with median synthetic-accessibility scores of 2.2-2.7 and median QED of 0.45-0.50. The AF3-predicted complexes of all nine leads, cealigned to their site anchors on the CFTR channel, show the designed candidates occupying the same pocket as the anchor on novel scaffolds (**Fig. 5c**).

**Table 4.** CFTR nine-candidate cohort and three clinical anchors, per molecule. (Trikafta sites A/B/C) and clinical anchors. iPTM / interface PAE from AlphaFold 3; Tan = ECFP4 Tanimoto to the site anchor; ChemFlag = reactive-group structural alert (blank = none). Full canonical SMILES shown.

| ID | SMILES | MW | logP | QED | Ro5 | Tan | iPTM | PAE | AF3 | Chem Flag |
| --- | --- | --- | --- | --- | --- | --- | --- | --- | --- | --- |
| **ivacaftor**<br>(anchor, site A) | CC(C)(C)C1=CC(=C(C=C1NC(=O)C2=CC3=CC=CC=C3C2=O)O)C(C)(C)C | 392 | 5.08 | 0.57 | 1 | 1.00<br>0 | 0.73 | 6.94 | WEAK | — |
| **tezacaftor**<br>(anchor, site B) | CC(C)(CO)C1=CC2=CC(=C(C=C2N1CC(CO)O)F)NC(=O)C3(CC3)C4=CC5=C(C(=C4)OC(O5)(F)F | 520 | 3.40 | 0.36 | 1 | 1.00<br>0 | 0.92 | 2.40 | MOD | — |
| **elexacaftor**<br>(anchor, site C) | CC1CC(N(C1)C2=C(C=CC(=N2)N3C=C(C(=N3)OCC(C)(C)C(F)(F)F)C(=O)NS(=O)(=O)C4=CN(N=C4C)C)(C)C | 598 | 4.02 | 0.41 | 1 | 1.00<br>0 | 0.73 | 7.93 | WEAK | — |
| **p10 (site C)** | CCCc1ccc(N=C(NC(=O)c2ccc(C)cc2)c2cc(OC)cc2)cc1 | 386 | 5.46 | 0.45 | 1 | 0.12<br>4 | 0.93 | 2.35 | MOD | — |
| **c671 (site C)** | CCCc1ccc(N=C(NC(=O)c2ccc(C)cc2F)c2ccc(OC)cc2)c(F)c1 | 422 | 5.74 | 0.41 | 1 | 0.14<br>3 | 0.92 | 2.59 | MOD | — |
| **L676 (site C)** | COc1ccc(C(=Nc2ccc(CCCO)cc2)NC(=O)c2ccc(C)cc2)cc1 | 388 | 4.44 | 0.50 | 0 | 0.12<br>1 | 0.77 | 7.01 | WEAK | — |
| **c11 (site B)** | CCc1ccc2nc(Nc3cc(C)cc(C#N)c3)n(-c3c(C)cccc3Br)c(=O)c2c1 | 487 | 5.94 | 0.46 | 1 | 0.09<br>5 | 0.93 | 2.28 | MOD | — |
| **c22 (site B)** | CCc1ccc2nc(Nc3cc(C)cc(C#N)c3)n(-c3cccc3Br)c(=O)c2c1C | 487 | 5.94 | 0.46 | 1 | 0.08<br>8 | 0.89 | 3.31 | MOD | — |
| **L716 (site B)** | Cc1cc(C#N)cc(Nc2nc3ccc(CCO)cc3c(=O)n2-c2c(C)cccc2Br)c1 | 489 | 4.92 | 0.46 | 0 | 0.11<br>1 | 0.88 | 3.57 | MOD | — |
| **c655 (site A)** | CCCc1ccc(N=C(NC(=O)c2ccc(C)cc2)c2cc(OC)cc2F)c(C#N)c1 | 414 | 5.48 | 0.50 | 1 | 0.11<br>7 | 0.69 | 7.36 | WEAK | — |
| **L1806 (site A)** | COc1ccc(C(=Nc2ccc(CCF)cc2C#N)NC(=O)c2ccc(C)cc2)cc1 | 418 | 4.90 | 0.50 | 0 | 0.11<br>2 | 0.69 | 7.17 | WEAK | — |
| **L1344 (site A)** | COCCc1ccc(Nc2cc(O)cc(CS)c2)n1 | 295 | 2.74 | 0.55 | 0 | 0.10<br>0 | 0.73 | 7.60 | WEAK | thiol |

To assess off-target liability, we cofolded the combinatorial candidate pool and the three clinical anchors against a CFTR off-target panel comprising four ABC-transporter paralogs from CFTR’s own gene family (ABCB1/P-gp, ABCC1/MRP1, ABCC4, ABCG2/BCRP) and the standard in vitro safety-pharmacology off-targets^45^ (the hERG potassium channel KCNH2 and the cytochrome-P450 enzymes CYP1A2, CYP2C9, CYP2D6, CYP3A4), classifying every cofold at the same decoy-calibrated thresholds (**Fig. S5**; **Table S4**). Across all nine off-target proteins, neither the designed candidates nor the three Trikafta anchors reached STRONG: both populations resolved at MODERATE confidence on the ABC transporters (designs median iPTM 0.78-0.85, interface PAE 3.0-4.6 Å; anchors median iPTM 0.77-0.81, interface PAE 3.9-6.4 Å) and on the CYP450 enzymes, and both resolved as WEAK on hERG (designs 99.4 % WEAK; all three anchors WEAK). The designs were therefore indistinguishable from the approved drugs across this panel. We interpret this as an AF3 co-folding ceiling for the large, polyspecific binding cavities of ABC transporters and CYP enzymes rather than as a STRONG-gate selectivity result, and we note that the hERG readout is subject to the AF3 hERG blind spot documented previously^5^ ; the panel establishes that the lead candidates present the same predicted off-target profile as the clinical anchors, not that they are free of these liabilities.

### Generative design and evolution of GLP1-family receptor agonists

The glucagon-like peptide-1 receptor (GLP1R) is a Class B G-protein-coupled receptor and the principal target of incretin therapies for type 2 diabetes and obesity. The most effective approved agents are injectable peptides. Semaglutide (Novo Nordisk), a GLP1R-selective agonist approved for type 2 diabetes and obesity, established the class^46^; tirzepatide (Eli Lilly), a dual agonist that additionally engages the glucose-dependent insulinotropic polypeptide receptor (GIPR), is approved for both indications and improved on semaglutide’s efficacy^47^; and retatrutide (Eli Lilly), a triple agonist that further adds the glucagon receptor (GCGR), reported still greater weight loss in a phase 2 trial^48^. The progression from single to dual to triple receptor engagement is the central pharmacological axis of the field: agonism at the two incretin paralogs GIPR and GCGR alongside GLP1R augments metabolic efficacy, so controlling the receptor-selectivity profile of an agonist is a primary design objective. The recent frontier is to recapitulate this pharmacology in orally available small molecules rather than peptides. Orforglipron (Eli Lilly), an oral GLP1R agonist, is in phase 3 development^49^; danuglipron (Pfizer), an oral GLP1R agonist, advanced to phase 2 before its clinical development was discontinued in 2025 following a case of potential drug-induced liver injury^50,51^; and TT-OAD2 is a preclinical small-molecule allosteric agonist used to resolve the GLP1R allosteric pharmacology structurally^52^. In the breadth screen, GLP1R returned 0 % STRONG under single-pass generation with 80 % of designs at MODERATE, the recoverable failure mode flagged earlier; this section develops the generative evolution recovery and shows that GenLoop produces small-molecule agonists whose receptor-selectivity profile is tunable across the single, dual, and triple tiers.

We ran a selectivity-aware instance of the four-step GenLoop cycle against a six-receptor panel comprising GLP1R, the two incretin paralogs GIPR and GCGR, and three additional Class B GPCRs as off-target safety controls (GLP2R, SCTR, PTHR1) (**Fig. 6a**). The target-conditioned decoder generated approximately 10,000 candidate molecules from the GLP1R sequence, of which about 4,900 were valid and unique. The encoder ranked these for predicted GLP1R binding and structural novelty, and the top-ranked candidates were cofolded against GLP1R in AF3. Consistent with the breadth screen (**Fig. 2d**), this single-pass pool bound GLP1R predominantly at MODERATE confidence and did not populate the STRONG tier; the five strongest and chemically distinct MODERATE-tier scaffolds became the AF3-verified parents for generative evolution.

**Figure 6.**
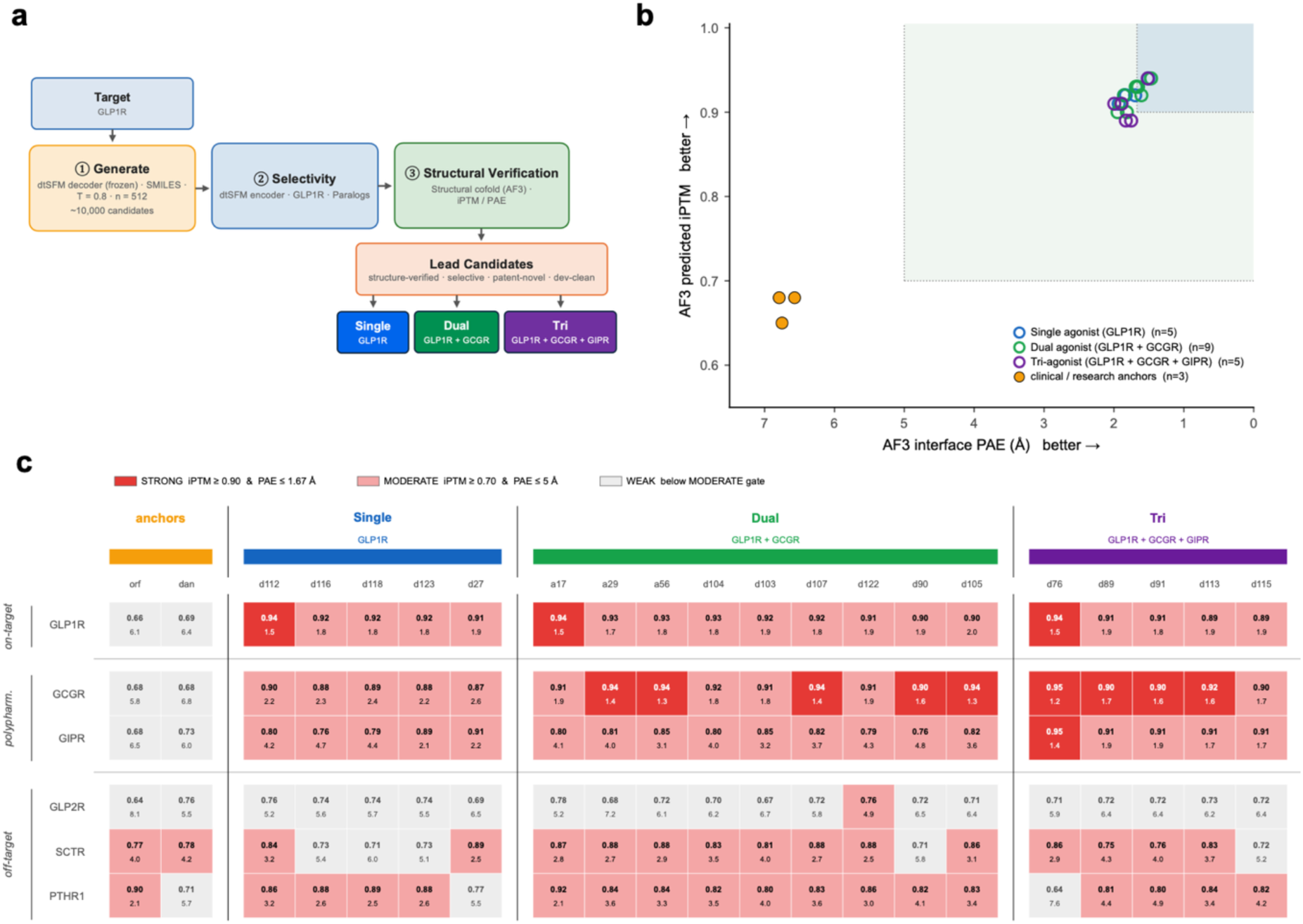
Selectivity-aware GenLoop for GLP1R. **(a)** GenLoop process schematic for GLP1R. Starting from ∼10,000 base-decoder designs, candidates are passed through the encoder selectivity panel (GLP1R target + 2 incretin paralogs GIPR / GCGR + 3 Class-B GPCR safety targets GLP2R / SCTR / PTHR1), then AF3-cofolded against all six receptors, then partitioned by selectivity tier: **PURE-GLP1R** (Single), **DUAL** (GLP1R + GCGR), **TRI** (GLP1R + GIPR + GCGR). Final cohort n = 22 (3 anchors + 5 Single + 9 Dual + 5 Tri). **(b)** AF3 verification, plotted as iPTM (vertical, better → top) vs interface PAE Å (horizontal, better → right). Designed candidates coloured by tier (Single = blue, Dual = green, Tri = purple); clinical / research-stage anchors (orforglipron, danuglipron, TT-OAD2) are gold filled circles. Shaded box marks the STRONG gate; designed candidates concentrate in the iPTM 0.85–0.95, PAE 1.5–2.5 Å region; anchors all WEAK (iPTM 0.65–0.68, PAE 6.6–6.8 Å). **(c)** AF3 selectivity profile across 6 Class-B GPCRs. Each row = one receptor (top → bottom: on-target GLP1R; polypharm partners GIPR, GCGR; off-target safety paralogs GLP2R, SCTR, PTHR1). Each column = one molecule, grouped left → right by tier (anchors gold, Single, Dual, Tri). Cell colour = AF3 class on that (molecule, receptor) cofold (STRONG = red, MODERATE = salmon, WEAK = gray); cell text = iPTM (top) / interface PAE Å (bottom).

From these five parents the library was built stepwise. A single-site bioisostere scan generated 73 variants, each cofolded against GLP1R in AF3; an interface-iPTM binding floor (iPTM ≥ 0.80) and a multi-objective Pareto filter over iPTM, logP, molecular weight, stereocentres, and synthetic accessibility identified 17 tolerated edits. Recombining only these tolerated edits produced a 124-member combinatorial library (53 single-extension, 39 double, and 32 triple substitutions). The single-site and combinatorial designs were then cofolded against all six receptors of the panel and classified at the decoy-calibrated joint thresholds, giving a six-receptor AF3 class vector per molecule; this selectivity-aware scoring both partitions the cohort by receptor profile and defines the off-site and paralog regressors used in the danuglipron-site refinement below. The selective designs partitioned into three tiers, Single (GLP1R only), Dual (GLP1R and GCGR), and Tri (GLP1R, GIPR, and GCGR), and the final cohort comprised 21 molecules: two clinical anchors (orforglipron, danuglipron) plus five Single, nine Dual (six combinatorial and three single-site), and five Tri designs (**Table 5**). Generative evolution recovered the STRONG tier that single-pass generation had missed: on the intended GLP1R target the designs concentrated at iPTM 0.85-0.95 and interface PAE 1.5-2.5 Å across all three tiers (tier-median iPTM 0.91-0.92, interface PAE 1.79-1.88 Å), reaching the STRONG gate, while both clinical anchors fell in the WEAK band (iPTM 0.69-0.72, interface PAE 5.2-6.4 Å; **Fig. 6b**). As in the CFTR campaign, we read the anchors as a same-fold reference rather than a STRONG gate: their low cofold scores reflect AF3’s limited confidence on these small-molecule Class B GPCR binding modes, so the designs’ STRONG cofolds are structurally confident poses for experimental testing, not evidence of superiority over the approved drugs.

**Table 5.**
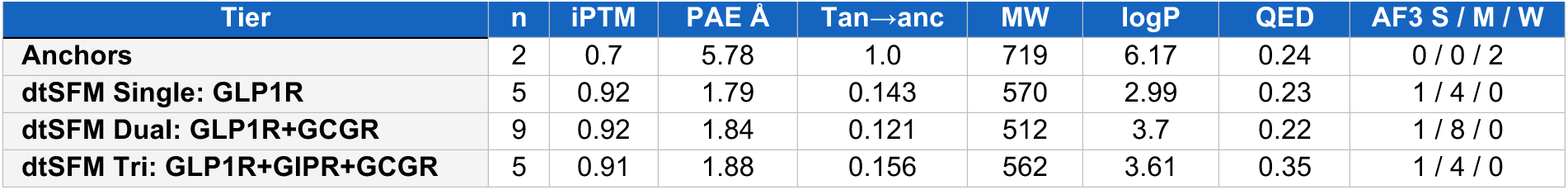
GLP1R candidate cohort, tier-level summary, mean values.

| Tier | n | iPTM | PAE Å | Tan→anc | MW | logP | QED | AF3 S / M / W |
| --- | --- | --- | --- | --- | --- | --- | --- | --- |
| **Anchors** | 2 | 0.7 | 5.78 | 1.0 | 719 | 6.17 | 0.24 | 0 / 0 / 2 |
| **dtSFM Single: GLP1R** | 5 | 0.92 | 1.79 | 0.143 | 570 | 2.99 | 0.23 | 1 / 4 / 0 |
| **dtSFM Dual: GLP1R+GCGR** | 9 | 0.92 | 1.84 | 0.121 | 512 | 3.7 | 0.22 | 1 / 8 / 0 |
| **dtSFM Tri: GLP1R+GIPR+GCGR** | 5 | 0.91 | 1.88 | 0.156 | 562 | 3.61 | 0.35 | 1 / 4 / 0 |

The selectivity result is the tiered receptor profile itself, a six-receptor classification heatmap (**Fig. 6c**; **Table 6**). We first confirmed that the panel is informative by reference to the clinical anchor: danuglipron, a GLP1R-selective agonist, was WEAK on all four non-GLP1R receptors GCGR, GIPR, GLP2R, and PTHR1, and MODERATE on SCTR, recapitulating its known receptor selectivity and establishing that the off-target safety paralogs resolve in the expected WEAK-to-MODERATE range rather than as an AF3 blind spot. Against this baseline, the designed cohort exhibited controllable polypharmacology: the on-target GLP1R row was STRONG or MODERATE across all 19 designs, the two incretin paralogs GIPR and GCGR turned STRONG only in the columns of the matching tier, and the three off-target safety paralogs GLP2R, SCTR, and PTHR1 were MODERATE or WEAK across all 19 designs. The loop therefore produced Single agonists that engage GLP1R alone, Dual agonists that add GCGR, and Tri agonists that add both GIPR and GCGR.

**Table 6.** GLP1R 19-candidate cohort and two clinical anchors, per molecule. iPTM / interface PAE from AlphaFold 3; Tan = ECFP4 Tanimoto to the site anchor; ChemFlag = reactive-group structural alert (blank = none). Full canonical SMILES shown.

| ID | SMILES | MW | logP | QED | Ro5 | Tan | iPTM | PAE | AF3 | Chem Flag |
| --- | --- | --- | --- | --- | --- | --- | --- | --- | --- | --- |
| **orforglipron (anchor)** | Cc1cc(-n2nc3c(c2-n2ccn(-c4ccc5c(cnn5C)c4F)c2=O)[C@H](C)N(C(=O)c2cc4cc([C@H]5CCOC(C)(C)C5)ccc4n2[C@@]2(c4noc(=O)[nH]4)C[C@@H]2C)CC3)cc(C)c1F | 883 | 7.44 | 0.17 | 2 | 1.000 | 0.72 | 5.15 | WEAK | — |
| **danuglipron (anchor)** | N#Cc1ccc(CO)c2ccccc(C3CCN(Cc4nc5ccc(C(=O)O)cc5n4C[C@@H]4CCO4)CC3)n2)c(F)c1 | 556 | 4.89 | 0.31 | 1 | 1.000 | 0.69 | 6.40 | WEAK | — |
| **d27 (Single)** | N#Cc1cc(F)cc(-c2nn(C(CN)c3ccc(C(=O)NCCC(=O)O)cc3)c3ccccc23)c1 | 472 | 3.47 | 0.36 | 0 | 0.120 | 0.91 | 1.92 | MOD | — |
| **d112 (Single)** | CC(C)(C)[C@H]1CC[C@]2(CC1)N=C(c1ccc(F)cc1)C(=O)N2[C@H](CO)c1ccc(C(=O)NCCC(=O)NO)cc1 | 553 | 3.75 | 0.29 | 1 | 0.174 | 0.94 | 1.52 | STRONG | hydroxamic |
| **d116 (Single)** | CC(C)(N)[C@H]1CC[C@]2(CC1)N=C(c1ccc(Cl)cc1)C(=O)N2[C@H](CO)c1ccc(C(=O)NCCC(=O)NO)cc1 | 570 | 2.95 | 0.23 | 1 | 0.143 | 0.92 | 1.79 | MOD | hydroxamic |
| **d118 (Single)** | CC(C)(O)[C@H]1CC[C@]2(CC1)N=C(c1ccc(Cl)cc1)C(=O)N2[C@H](CO)c1ccc(C(=O)NCCC(=O)NO)cc1 | 571 | 2.99 | 0.23 | 1 | 0.135 | 0.92 | 1.82 | MOD | hydroxamic |
| **d123 (Single)** | CC(N)(F)[C@H]1CC[C@]2(CC1)N=C(c1ccc(Cl)cc1)C(=O)N2[C@H](CO)c1ccc(C(=O)NCCC(=O)NO)cc1 | 574 | 2.86 | 0.18 | 1 | 0.159 | 0.92 | 1.75 | MOD | hydroxamic ; gem-N,F |
| **d76 (Tri)** | N#Cc1cc(Cl)cc(-c2nn(C(CN)c3ccc(C(=O)NCCc4nnn[nH]4)cc3)c3ccccc23)c1 | 512 | 3.26 | 0.29 | 1 | 0.088 | 0.94 | 1.48 | STRONG | — |
| **d91 (Tri)** | CC(C)(C)[C@H]1CC[C@]2(CC1)N=C(c1ccc(F)cc1)C(=O)N2[C@H](CO)c1ccc(C(=O)NCCc2nnn[nH]2)cc1 | 562 | 3.61 | 0.38 | 1 | 0.160 | 0.91 | 1.76 | MOD | — |
| **d89 (Tri)** | CC(C)(C)[C@H]1CC[C@]2(CC1)N=C(c1ccc(F)cc1)C(=O)N2[C@H](CO)c1ccc(C(=O)NCCc2nnn[nH]2)cc1 | 562 | 3.61 | 0.38 | 1 | 0.156 | 0.91 | 1.90 | MOD | — |
| **d113 (Tri)** | CC(C)(F)[C@H]1CC[C@]2(CC1)N=C(c1ccc(Cl)cc1)C(=O)N2[C@H](CO)c1ccc(C(=O)NCCc2nnn[nH]2)cc1 | 582 | 3.83 | 0.35 | 1 | 0.145 | 0.89 | 1.90 | MOD | — |
| **d115 (Tri)** | CC(C)(F)[C@H]1CC[C@]2(CC1)N=C(c1ccc(Cl)cc1)C(=O)N2[C@H](CO)c1ccc(C(=O)NCCc2nnn[nH]2)cc1 | 582 | 3.83 | 0.35 | 1 | 0.158 | 0.89 | 1.88 | MOD | — |
| **d104 (Dual)** | N#Cc1cc(Cl)cc(-c2nn(C(CO)c3ccc(C(=O)NCCC(=O)NO)cc3)c3ccccc23)c1 | 504 | 3.44 | 0.21 | 1 | 0.102 | 0.93 | 1.75 | MOD | hydroxamic |
| **d103 (Dual)** | N#Cc1cc(Cl)cc(-c2nn(C(CN)c3ccc(C(=O)NCCC(=O)NO)cc3)c3ccccc23)c1 | 503 | 3.40 | 0.21 | 1 | 0.101 | 0.92 | 1.85 | MOD | hydroxamic |
| **d107 (Dual)** | O=C(CCNC(=O)c1ccc(C(CO)n2nc(-c3cc(F)cc(Cl)c3)c3ccccc32)cc1)NO | 497 | 3.70 | 0.22 | 0 | 0.122 | 0.92 | 1.84 | MOD | hydroxamic |
| **d105 (Dual)** | NCC(c1ccc(C(=O)NCCC(=O)NO)cc1)n1nc(-c2cc(F)cc(Cl)c2)c2ccccc21 | 496 | 3.67 | 0.22 | 0 | 0.121 | 0.90 | 1.97 | MOD | hydroxamic |
| **d90 (Dual)** | CC(C)(F)[C@H]1CC[C@]2(CC1)N=C(c1cccc(F)c1)C(=O)N2[C@H](CO)c1ccc(C(=O)NCCC2nnn[nH]2)cc1 | 566 | 3.31 | 0.36 | 1 | 0.177 | 0.90 | 1.94 | MOD | — |
| **a56 (Dual)** | NCC(c1ccc(C(=O)NCCC(=O)NO)cc1)n1nc(-c2cc(Cl)cc(Cl)c2)c2cccc21 | 512 | 4.18 | 0.21 | 1 | 0.098 | 0.93 | 1.75 | MOD | hydroxamic |
| **a29 (Dual)** | O=C(CCNC(=O)c1ccc(C(CO)n2nc(-c3cc(Cl)cc(Cl)c3)cccc32)cc1)NO | 513 | 4.22 | 0.21 | 1 | 0.099 | 0.93 | 1.71 | MOD | hydroxamic |
| **a17 (Dual)** | CC(C)(C)[C@H]1CC[C@]2(CC1)N=C(c1ccc(Cl)cc1)C(=O)N2[C@H](CO)c1ccc(C(=O)NCCC(=O)NO)cc1 | 569 | 4.26 | 0.28 | 1 | 0.136 | 0.94 | 1.50 | STRONG | hydroxamic |
| **d122 (Dual)** | CC(C)(F)[C@H]1CC[C@]2(CC1)N=C(c1cccc(Cl)c1)C(=O)N2[C@H](CO)c1ccc(C(=O)NCCC(=O)NO)cc1 | 573 | 3.96 | 0.27 | 1 | 0.175 | 0.91 | 1.86 | MOD | hydroxamic |

To engineer molecules occupying the danuglipron central pocket specifically, we applied a round of generative evolution refinement with a danuglipron-site LoRA, trained as in the CFTR site-C campaign and followed by a reseed after an initial hydroxamic-acid reactive-group flag. The refined cohort comprised 23 candidates, all of which matched the danuglipron pocket by gemmi canonical-frame site assignment (23 of 23 SITE_MATCH), at median iPTM 0.89 and interface PAE 1.95 Å (**Table 6**). The reseed reduced but did not eliminate this motif: 12 of the 19 tabulated leads retain a terminal hydroxamic-acid bioisostere of the anchor carboxylate, a Brenk structural alert, now flagged in **Table 6**, while 7 are fully Brenk-clean, having converged on the carboxylic-acid or tetrazole bioisostere present elsewhere in the same cohort; the hydroxamic group is a single-step swap that defines the next optimisation round of GenLoop. The AF3-predicted complexes of the cohort, overlaid with the danuglipron anchor across the GLP1R, GIPR, and GCGR pockets and organised by tier, occupied the central pocket (**Fig. 7**). The campaign was bounded by one out-of-distribution limitation: the Gs-biased allosteric pocket engaged by orforglipron, approximately 35 Å from the danuglipron central pocket, was not reached, as an analogously trained orforglipron-site LoRA failed to place candidates at the allosteric site, a structural context underrepresented in the dtSFM training corpus.

**Figure 7.**
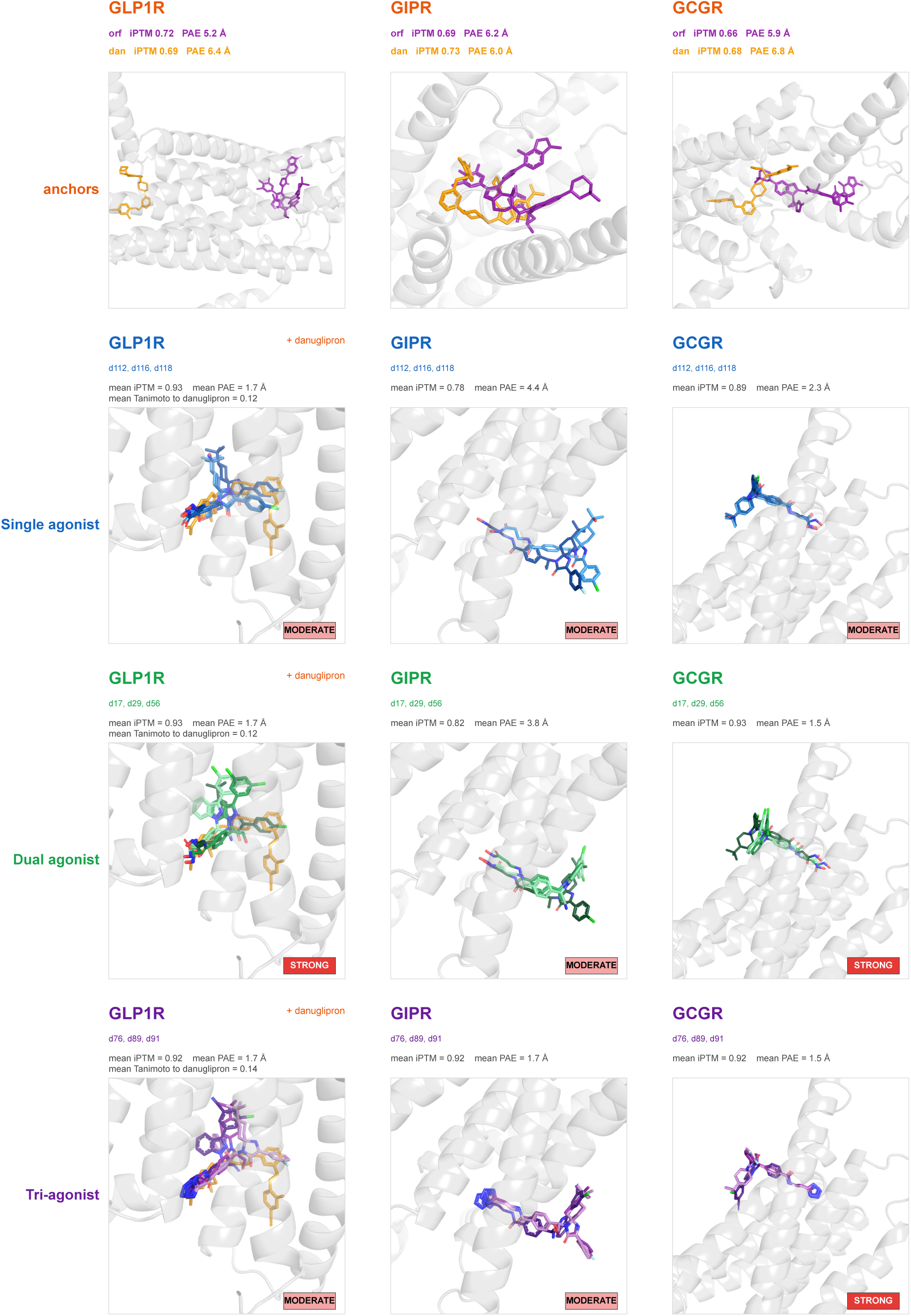
Multi-overlay AF3 gallery for the dtSFM-designed GLP1-family targets. Each cell shows the AF3-predicted complex of three designed candidates (sticks) overlaid in a single per-receptor camera so candidates are directly comparable across the receptor row, plus the danuglipron clinical anchor (gold) shown where it occupies the GLP1R column (anchor row only renders the three anchors directly). The receptor backbone is rendered as semi-transparent gray cartoon, identical camera per receptor. Cells are labelled with: receptor name (coloured by row tier); design identifiers (dNNNN, three per cell); mean AF3 iPTM and mean interface PAE Å across the three overlays; for GLP1R cells, mean Tanimoto to danuglipron. Each cell carries a class chip (STRONG = red, MODERATE = salmon, WEAK = gray) for the tier-mean class on that receptor.

As an orthogonal cross-model check, the GLP1R and CFTR cohort and the clinical anchors were independently cofolded with Boltz-2; the AF3 and Boltz-2 structural-confidence metrics agree across the designed cohort (**Fig. S7**).

## DISCUSSION

The base decoder of dtSFM generates target-conditioned chemistry in a single pass^5^, but a single pass is bounded: per-target structural yield ranged from complete to absent across the breadth panel, and selectivity, multi-site depth, and tunable polypharmacology lie outside its scope. GenLoop closes the cycle, and these capabilities emerge from iteration rather than from a larger model. The loop is directed evolution in the protein-engineering sense^26,27^, a library generated, selected for on-target function, counter-selected against off-target binding, and advanced through multi-parameter optimisation, with the difference that its library is sampled from the sequence-native dtSFM decoder by computational thermodynamic selection, its selector is an independent structural verifier (AF3), and its multi-parameter optimisation is a stack of cheminformatics developability gates. The discriminating power of GenLoop comes from pairing the dtSFM encoder, which scores a candidate’s compatibility with the target, against a structural verifier (AF3) orthogonal to that scoring axis, so that the candidates carried forward are those on which two AI models sharing no representation independently agree.

This mirrors how chemical matter advances in practice: a discovery campaign does not advance its first-pass hits but carries a diverse, liability-bearing pool into lead optimisation. The breadth screen is that hit-generation step — a third of the anchor-confident designs were already free of reactive-group flags, and the remainder concentrate the decoder’s terminal-motif artefacts that an optimisation phase exists to remove. The CFTR and GLP1-family campaigns are that optimisation phase rendered in silico as generative evolution: dtSFM proposes site-directed chemical variations and rescreens them in the loop — the small-molecule analogue of mutate-and-select directed evolution — recovering the selectivity, multi-site depth, and tunable polypharmacology that no single pass provides. Optimisation is itself iterative: even the GLP1-family cohort retains a terminal hydroxamic-acid bioisostere of the anchor carboxylate in 12 of its 19 designs (**Table 6)**, a Brenk structural alert and a single-step swap to the carboxylic-acid or tetrazole present among the clean designs, so the loop yields advanced leads rather than finished molecules.

GenLoop produced lead candidates across mechanistically distinct settings. In the breadth screen, candidates reached clinical-anchor structural confidence across kinase, nuclear-receptor, complement-protease, protein-interaction, and G-protein-coupled-receptor targets, occupying the same pockets as approved drugs with scaffolds novel by Tanimoto. On CFTR, the loop delivered nine lead candidates across the three orthogonal Trikafta sites at Tanimoto 0.09 to 0.14 to their clinical anchors, with a round of generative evolution enriching occupancy of the deepest site 4.8-fold. On the GLP1-family receptors, the loop recovered a target that single-pass generation had left at zero structurally confident designs, producing a cohort whose receptor-selectivity profile is tunable across the single, dual, and triple incretin tiers, all matched to the central agonist pocket and engineered into a favourable lipophilicity range. Relative to the two clinical anchors, the dtSFM cohort sidesteps both development liabilities — the Hy’s-law liver toxicity that halted danuglipron’s Phase 2b and the high lipophilicity (logP > 5) of orforglipron — at a median design logP of 3.6, while matching the danuglipron central pocket and remaining structurally novel to both (ECFP4 Tanimoto ≤ 0.18). Across all three settings the candidates are simultaneously structure-verified against the intended target, resolved against paralogs and off-targets, novel by scaffold, and filtered for synthesizability and drug-likeness, which is to say they are multi-parameter-optimised cohorts rather than single-property hit lists.

The structural confidence that AF3 assigns to dtSFM-generated molecules, consistent across mechanistically distinct settings, has a thermodynamic explanation. An SFM does not approximate an unknown function; it estimates the parameters of a known one: the convergence equation dictates that the softmax attention mechanism of transformers that scores compatibility is mathematically isomorphic to the Boltzmann distribution that governs molecular binding at thermal equilibrium^1,2^. dtSFM therefore performs computational thermodynamics: its embedding is a free-energy-like compatibility between a drug and a target, and its decoder samples chemistry from a learned, target-conditioned distribution over chemical space in which thermodynamically compatible molecules are the high-probability draws. Generation is thus thermodynamic selection rather than pattern completion, which is why a sequence-native model that never constructs a three-dimensional coordinate produces molecules that an independent structure-prediction model places in the correct pocket: the two computations are estimating the same physical quantity by different routes, and their agreement is corroboration from distinct physical principles. GenLoop closes the generative loop around dtSFM, bringing the library, selection, and multi-parameter optimisation of directed evolution to a model that computes molecular recognition as thermodynamic selection, and so turns a single-pass specificity foundation model into an iterative engine for sequence-native drug design.

Every result reported here is a structural prediction, and the GenLoop candidate cohorts, model weights, decoder, refinement recipe, verification protocol, and developability gates are released in full so that any laboratory can run the loop, extend it with proprietary data, and carry its designs forward. The platform is bounded by the coverage of its training corpus and by the per-target scope of its generative evolution refinement, but its decisive limitation is that target engagement is computationally predicted and not yet experimentally measured. Wet-lab validation is the step that closes the generative loop in the laboratory rather than in silico, and it is the essential next test of the designs reported here.

## METHODS

### Compute infrastructure

Decoder sampling, encoder reranking, generative evolution refinement, and all downstream cheminformatics processing were performed on Euler, ETH Zurich’s academic HPC cluster, with GPU jobs running on single NVIDIA A100 40 GB or comparable cards. The LoRA adapters were lightweight enough to train on a single consumer-grade GPU: each adapter completed 1,000 optimisation steps in approximately three minutes on an NVIDIA TITAN RTX, so that the generative evolution refinement step does not require HPC resources. AF3 cofolding at scale, comprising more than 15,000 individual cofolds across the breadth, selectivity, and depth campaigns, was performed on the ALPS supercomputer at the Swiss National Supercomputing Centre (CSCS), whose Clariden compute nodes are built on NVIDIA Grace Hopper Superchip (GH200) modules pairing a Hopper-class H100 GPU with a Grace ARM CPU.

### dtSFM base model

GenLoop operates on dtSFM version 3, the drug-target specificity foundation model whose architecture, training corpus, and four-head validation are described in full in the companion paper^5^; only the elements required to reproduce the loop are restated here. The model couples two frozen pretrained encoders, MoLFormer-XL^53^ on the chemical side, which embeds a canonical SMILES string into a 768-dimensional vector, and ESM-2-650M ^54^ on the protein side, which embeds a target sequence into a 1,280-dimensional per-residue representation, through a trainable cross-attention encoder of 14.4 million parameters that emits four supervised heads (retrieval, interface, residue contact, and affinity). Target-conditioned generation uses the cross-attentive autoregressive SMILES decoder of the same model, a 27-million-parameter decoder-only Transformer with masked self-attention over generated tokens and cross-attention over the projected target features, which is the first generative instance of the SFM architecture^2^. All GenLoop campaigns used the production encoder checkpoint epoch_010 and the decoder checkpoint decoder_v0.2_step50K, both loaded with frozen weights; the cheminformatics, reranking, and refinement operations described below act on these fixed checkpoints without retraining the base model.

### Target-conditioned decoder sampling

For each target, the autoregressive SMILES decoder was conditioned on the target’s ESM-2 per-residue features and sampled with multinomial decoding at temperature T = 0.8, with end-of-sequence termination and a maximum generation length of 120 tokens. The default per-target budget was n = 512 raw samples for the breadth screen; the depth campaigns sampled larger pools, including approximately 10,000 samples per target for the GLP1R polypharmacology campaign and the CFTR site campaigns. Generated token sequences were detokenised to SMILES strings and parsed with RDKit^31^; sequences that failed to parse to a valid molecule were discarded, and the surviving molecules were canonicalised and deduplicated on their canonical SMILES so that each unique structure entered the loop once.

### Encoder reranking

Each unique candidate was scored against the intended target by the dtSFM retrieval head, which computes the cosine similarity between the drug and target global embeddings in the shared 512-dimensional space. Candidates were ranked by this drug-target cosine, and the top 20 % of each per-target pool were forwarded to structural verification, a triage that concentrates the pool on target-relevant chemistry without collapsing its scaffold novelty. For the selectivity-aware campaigns, the rerank was generalised from a single target to a receptor panel: each candidate was scored against every member of the panel, and candidates were prioritised by their full cross-receptor cosine profile rather than by on-target cosine alone, so that the molecules forwarded to cofolding spanned the intended selectivity tiers. The encoder cosine is a relevance triage and is statistically independent of the downstream AF3 structural confidence^5^ ; the two are applied as orthogonal filters.

### Structural verification and site assignment

Reranked candidates were verified by cofolding each (drug, target) pair in AF3 ^21^, run from its licensed model weights inside an Enroot container on the ALPS GH200 nodes. Multiple-sequence alignments were pre-generated for every target with MMseqs2^55^ in place of the default jackhmmer pipeline, cleaned to the AF3 input schema, and injected as the unpaired MSA, with the paired MSA left empty and templates disabled; each cofold was run as a single model with five diffusion seeds. From each cofold we extracted the interface predicted TM score (iPTM) and the minimum interface predicted aligned error (interface PAE) from the AF3 confidence output.

Each cofold was assigned a binding-confidence class from its (iPTM, interface PAE) pair against decoy-calibrated joint thresholds: STRONG (iPTM >= 0.90 and interface PAE <= 1.67 A), MODERATE (iPTM >= 0.70 and interface PAE <= 5.00 A), and WEAK otherwise. The thresholds were calibrated on a panel of n = 24 negative-control molecules, drug-like SMILES with no documented relationship to any target in the screen, cofolded against each of the 12 breadth targets: 0 % of decoy cofolds reached STRONG and 33 % reached MODERATE, so the STRONG gate defines specific binding (0 of 24 decoys reached STRONG, a one-sided 95 % upper bound of ≈12 %) and the MODERATE gate is reported as ambiguous (within the AF3 noise floor). The thresholds and the classifier are defined once in a single module (genloop_defs) and imported by every figure, table, and analysis script so that the cutoffs do not drift.

For the depth campaigns, where multiple distinct pockets exist on one target, each cofolded ligand was assigned to a binding site by canonical-frame geometry using gemmi^44^. The predicted complex was superposed onto a common reference frame for the target, the centroid of the designed ligand was computed, and its distance to the centroid of each clinical anchor’s pocket was measured; a candidate was recorded as a strict site match (SITE_MATCH) to a given site when its ligand centroid lay within 8 A of that site’s anchor pocket. This procedure assigned the CFTR candidates to the ivacaftor (site A), tezacaftor (site B), and elexacaftor (site C) pockets, and the GLP1R candidates to the danuglipron central pocket versus the orforglipron allosteric pocket. Structural overlays were rendered in PyMOL by superposing each candidate’s predicted complex onto the clinical anchor’s with the cealign algorithm under a single fixed per-target or per-site pocket camera (designed ligand blue, anchor gold, target as semi-transparent grey cartoon), and two-dimensional chemical structures were drawn with RDKit.

### Library design (bioisostere diversification)

For the generative evolution depth campaigns, candidate libraries were built from AF3-verified parent scaffolds with a curated RDKit SMARTS reaction library of about 80 bioisostere transforms across seven categories (lipophilicity reduction, halogen swap, scaffold hop, functional-group swap, ring saturation, chain shortening, and bioisosteric replacement; for example, carboxylic acid to tetrazole, oxadiazolone, or sulfonamide; amide to oxadiazole; CF₃ to CN or CHF₂). Edits were introduced singly (a single-site scan applying every transform at every matching position) and in combination (single-extension, double-, and sampled, capped triple-substitution variants), stepwise or in parallel depending on the campaign; all variants were sanitized, canonicalized, deduplicated, and developability-filtered, and were scored and selected with the dtSFM encoder cosine and AF3 cofolding. GLP1R used a stepwise single-site-then-combinatorial design: from 5 parents, the single-site scan produced 73 variants, each AF3-cofolded against GLP1R, gated at iPTM ≥ 0.80 and ranked by a five-objective Pareto filter (maximize interface iPTM; minimize logP, molecular weight, stereocentre count, synthetic-accessibility score), yielding 17 tolerated edits; recombining only these gave 124 combinatorial hybrids (53 single-extension, 39 double, 32 triple), cofolded against the six-receptor incretin panel and partitioned into selectivity tiers; the danuglipron-site refinement expanded 7 seed scaffolds into a 287-variant bioisostere pool (LoRA training set: 287 positives and 287 paralog-regressor negatives), yielding the 23-candidate cohort. CFTR used a combinatorial design: an 845-candidate combinatorial round built on the AF3-verified per-site shortlists was cofolded across CFTR and the 9 paralogs (about 8,450 cofolds) and scored by the dtSFM encoder cosine to CFTR, producing the 929 site-resolved verified cofolds; a site-C LoRA round (344 training molecules: 61 site-C positives and 283 off-site negatives) raised strict site-C occupancy from 7.5 % to 36.0 %.

### LoRA directed-evolution refinement

The directed-evolution step refined the decoder toward a specific binding site by low-rank adaptation (LoRA)^29^, implemented with the PEFT library. LoRA adapters were attached to the attention projections of the frozen 27-million-parameter base decoder, adding 688,000 trainable parameters, 2.5 % of the base, while all base weights remained fixed. Each adapter was trained on a site-resolved set of structurally verified molecules drawn from the preceding cofold round: positives were candidates that AF3 placed at the intended site by gemmi site assignment, and negatives were candidates placed at off-target sites or paralog pockets. Training minimised an unlikelihood objective^30^ that combined a standard likelihood term raising the probability of the positive SMILES with an unlikelihood term lowering the probability of the negative SMILES, so that refinement both concentrated generation on the intended pocket and pushed it away from the off-site chemistry that the verification round had surfaced. Adapters were trained for 1,000 steps, after which the adapted decoder was resampled under the same settings as the base decoder and the resulting candidates re-entered the loop. For the CFTR site-C campaign the positive and negative sets comprised 61 and 283 molecules respectively; the GLP1R danuglipron-site campaign used 287 positives and 287 negatives, and was followed by a reseed after an initial round flagged a reactive hydroxamic-acid group.

### Cheminformatics developability filtering

Candidates that passed structural verification were filtered for developability with a stack of cheminformatics gates computed in RDKit^31^. Reactive and unstable chemistries were removed with SMARTS substructure queries^32^ drawn from the Brenk panic-alert set^33^, and pan-assay-interference compounds were removed with the PAINS substructure filters^34^. Drug-likeness and physicochemical properties were assessed with the Lipinski Rule-of-Five^9^, the quantitative estimate of drug-likeness (QED)^35^, and the synthetic-accessibility score^36^, and the synthesizability gate was tuned against the make-on-demand chemistry of a commercial vendor library so that surviving candidates fall within tractable synthesis space. Scaffold novelty and structural novelty were quantified by Bemis-Murcko scaffold decomposition^37^ and by Morgan extended-connectivity fingerprint (ECFP4, radius 2, 2,048 bits) Tanimoto similarity^38^ of each candidate to the clinical anchors of its campaign, with a Tanimoto below 0.40 to the nearest anchor taken as the conventional threshold for a structurally novel scaffold.

### Paralog and off-target panels

Selectivity was assessed by cofolding designed candidates and their clinical anchors against curated panels of related proteins, scored and classified identically to the on-target cofolds. For the breadth screen, paralog panels were assembled for the 9 of 12 targets with structurally characterised families (ALK, BTK, CFB, GCGR, GIPR, GLP1R, MEN1, THRB, TYK2), spanning 3 to 6 paralogs each for 41 paralogs in total; the remaining three targets (JAK3, NLRP3, SHP2) were left without panels and are disclosed as a coverage gap. The CFTR depth campaign was screened against an off-target panel of four ABC-transporter paralogs (ABCB1/P-glycoprotein, ABCC1/MRP1, ABCC4, ABCG2/BCRP) and a five-member safety panel comprising the hERG potassium channel (KCNH2) and the cytochrome-P450 enzymes CYP1A2, CYP2C9, CYP2D6, and CYP3A4. The GLP1R polypharmacology campaign was screened against a six-receptor panel of the intended target GLP1R, the two incretin paralogs GIPR and GCGR, and three additional Class B G-protein-coupled receptors as off-target controls (GLP2R, SCTR, PTHR1). In every campaign the clinical anchors were cofolded through the same panel as an internal control on the informativeness of the AF3 readout, since a panel on which the approved drug shows its known selectivity profile validates the panel, whereas one on which the anchor and the designs are indistinguishable indicates a co-folding ceiling rather than a selectivity result.

### Breadth screen

The breadth screen applied the loop to 12 targets spanning kinase (ALK, BTK, JAK3, TYK2), nuclear-receptor (THRB), complement-protease (CFB), protein-interaction (MEN1), phosphatase (SHP2), inflammasome (NLRP3), and Class B GPCR (GCGR, GIPR, GLP1R) modalities, each with a clinical or research-stage anchor for reference. For each target, 512 raw candidates were sampled from the base decoder, filtered for RDKit validity and uniqueness, and reranked by encoder cosine, and a representative top-ranked set was cofolded in AF3, yielding 238 designed-candidate cofolds.

### CFTR depth campaign

The CFTR depth campaign ran the loop using the corresponding clinical molecule as the site anchor: ivacaftor (potentiator, site A), tezacaftor (Type II corrector, site B), and elexacaftor (Type III corrector, site C). Candidates were generated, encoder-reranked, cofolded against CFTR, and assigned to sites by gemmi, producing 929 site-resolved cofolds (51 at site A, 733 at site B, 145 at site C). Because the initial combinatorial round under-sampled the elexacaftor pocket (7.5 % strict site-C SITE_MATCH), a site-C LoRA refinement was applied, raising strict site-C SITE_MATCH to 36.0 % (**Fig. S6**). Three lead candidates per site were selected after developability filtering, and the full pool plus the three anchors were screened against the CFTR off-target panel of four ABC transporters and five ADMET safety proteins (**Fig. S5**; **Table S4**).

### GLP1R polypharmacology campaign

The GLP1R polypharmacology campaign ran the selectivity-aware loop against the six-receptor panel, using orforglipron and danuglipron as anchors. Approximately 10,000 base-decoder candidates were reranked against the full panel, cofolded against all six receptors, and partitioned by their AF3 selectivity profile into Single (GLP1R only), Dual (GLP1R and GCGR), and Tri (GLP1R, GIPR, and GCGR) tiers, yielding a 22-candidate cohort of three anchors and 19 designs (**Fig. 6**; **Tables 5** and **6**). A danuglipron-site LoRA refinement, followed by a reseed, produced a 23-candidate central-pocket cohort (23 of 23 SITE_MATCH; **Fig. 7**; **Table 6**). An orforglipron-site LoRA trained by the same procedure did not yield candidates that AF3 placed at the allosteric pocket, which is out of distribution for the base model.

### Orthogonal AI verification

#### Structural verification

Every structural prediction in this work was checked by AlphaFold-3 (AF3) as the primary orthogonal verifier. AF3 shares no architecture, training data, or learned representation with dtSFM, and on the candidate cohorts the dtSFM encoder cosine is essentially uncorrelated with AF3 interface confidence (Pearson r ≈ 0); structural agreement therefore constitutes independent corroboration rather than circular confirmation^5^. A subset of predictions was additionally cross-checked with Boltz-2, an independent implementation of the same cofolding paradigm (**Fig. S7**).

#### Coding audit

Every quantitative claim in this paper was independently re-derived by a second AI coding assistant (Codex, OpenAI) acting as an orthogonal auditor. Operating in a clean session with access only to the artifacts committed to the public repository — and no access to the development sessions that produced them — the auditor recomputed each claim from its source data and graded it pass, partial, or fail. The audit closed green across the full 34-claim set at 25 pass / 9 partial / 0 fail, with no numerical claim found to be in error (the partials are documented bookkeeping items, not defects). The complete audit record, the claim-to-file map, and all verification scripts are provided in the code repository (https://github.com/Reddy-BIIE-ETHZ/GenLoop) and the archived data deposit (https://doi.org/10.5281/zenodo.20626910).

### Vibe Coding Starter Prompt — running GenLoop on your target

GenLoop is designed for direct use by experimental biologists with no coding background. The block below is a single prompt: paste it into an AI Coding Agent such as Claude Code (Anthropic) along with the published GenLoop PDF. Claude Code will pull the repository and the dtSFM model weights, install the environment, and walk you through your campaign in plain language. The only fields you must fill in are your name and the campaign checkbox; every other field is optional, since Claude Code can fetch the target sequence from UniProt, identify the closest approved binder, assemble a paralog panel, and choose default hyperparameters by itself when those fields are left blank.

> *I am [YOUR NAME], a [ROLE — optional] at [INSTITUTION — optional]*
>
> *I am an experimental biologist with NO coding experience. Please run in operating mode — explain everything in plain language as you go, and issue commands only when one is genuinely necessary (which should be almost never)*.
>
> *I want to run a GenLoop campaign to design small-molecule candidates for:*
>
> *[] de novo lead generation — for which target?*
>
> *[optional: gene symbol or UniProt ID; if blank you may ask me or look it up yourself]*
>
> *[] multi-site depth — a target with more than one druggable pocket, designing leads at each site*
>
> *[optional: gene symbol or UniProt ID, and the sites or anchor drugs if you know them]*
>
> *[] selectivity-aware / polypharmacology — tuning engagement across a target family*
>
> *[optional: the intended target plus the paralogs to engage and the paralogs to avoid]*
>
> *[] directed-evolution refinement — focusing generation on one specific binding site*
>
> *[optional: the target and the site or anchor drug that defines the pocket]*
>
> *Optional context (leave blank if you don’t know — you can fetch it yourself from UniProt, ChEMBL, BindingDB, DrugBank, or the PDB):*
>
> - *target protein sequence (FASTA)*
>
> - *UniProt accession*
>
> - *closest clinically approved or research-stage binder, if any*
>
> - *the binding sites or paralogs that matter for selectivity*
>
> - *any in-house wet-lab labels I have for this target*
>
> *Pull the GenLoop repository and the dtSFM model weights from the URLs in the attached paper (Methods, “Compute infrastructure”). Install the runtime environment. Read the relevant Methods subsections — most likely “Target-conditioned decoder sampling”, “Encoder reranking”, “Structural verification and site assignment”, “LoRA directed-evolution refinement”, and “Cheminformatics developability filtering”. Then run the loop on my target: generate candidates, rerank them, verify them structurally, filter them for developability, and, if I asked for depth, selectivity, or refinement, run the corresponding panel and LoRA step. For the structural-verification step, use AF3 if I have access to its licensed weights; otherwise use the open-source Boltz-2 cofolding model, which produces the same interface confidence metrics, and tell me which one you used. If you use Boltz-2, re-calibrate the STRONG and MODERATE thresholds on the decoy panel for that model first, since the published thresholds were calibrated on AF3. Show me the resulting lead cohort in plain language with brief plots or tables, distinguish clearly between what is predicted and what would require wet-lab measurement, and ask me clarifying questions only when you genuinely need something I have not provided*.

The structural-verification step uses a protein-ligand cofolding model to score each candidate. AF3 requires access to its licensed model weights; where that access is unavailable, the open-source Boltz-2 model^23^ can be used in its place, producing the same interface confidence metrics, so the loop runs end to end without licensed software. The decoy-calibrated STRONG and MODERATE thresholds reported here were derived on AF3; for a campaign run on Boltz-2, it is advised to re-derive thresholds on the decoy and anchor panel for before classifying candidates.

## Data and code availability

GenLoop builds on the pre-trained dtSFM v3 model (weights: https://huggingface.co/SFM-BIIE-ETHZ/dtSFM-v3). All code, data tables, figures, and orthogonal audit records are on GitHub: https://github.com/Reddy-BIIE-ETHZ/GenLoop. The designed molecules are available as a Hugging Face dataset (https://huggingface.co/datasets/SFM-BIIE-ETHZ/GenLoop) and their AlphaFold 3 cofold structures on Zenodo (https://doi.org/10.5281/zenodo.20626910). All molecules disclosed here are dedicated to the public domain and free for any use; the dtSFM model and GenLoop method are released for research under the SFM Research Preview License (see LICENSE.md in the GitHub repository).

## Acknowledgments

ETH Zurich Scientific IT Services for High Performance Computing and Swiss National Supercomputing Centre (CSCS) provided computing resources and excellent support. Thank you to Riyaz Khan for Boltz-2 cofolding analysis and additional feedback.

## Supplementary Material

**Figure S1.**
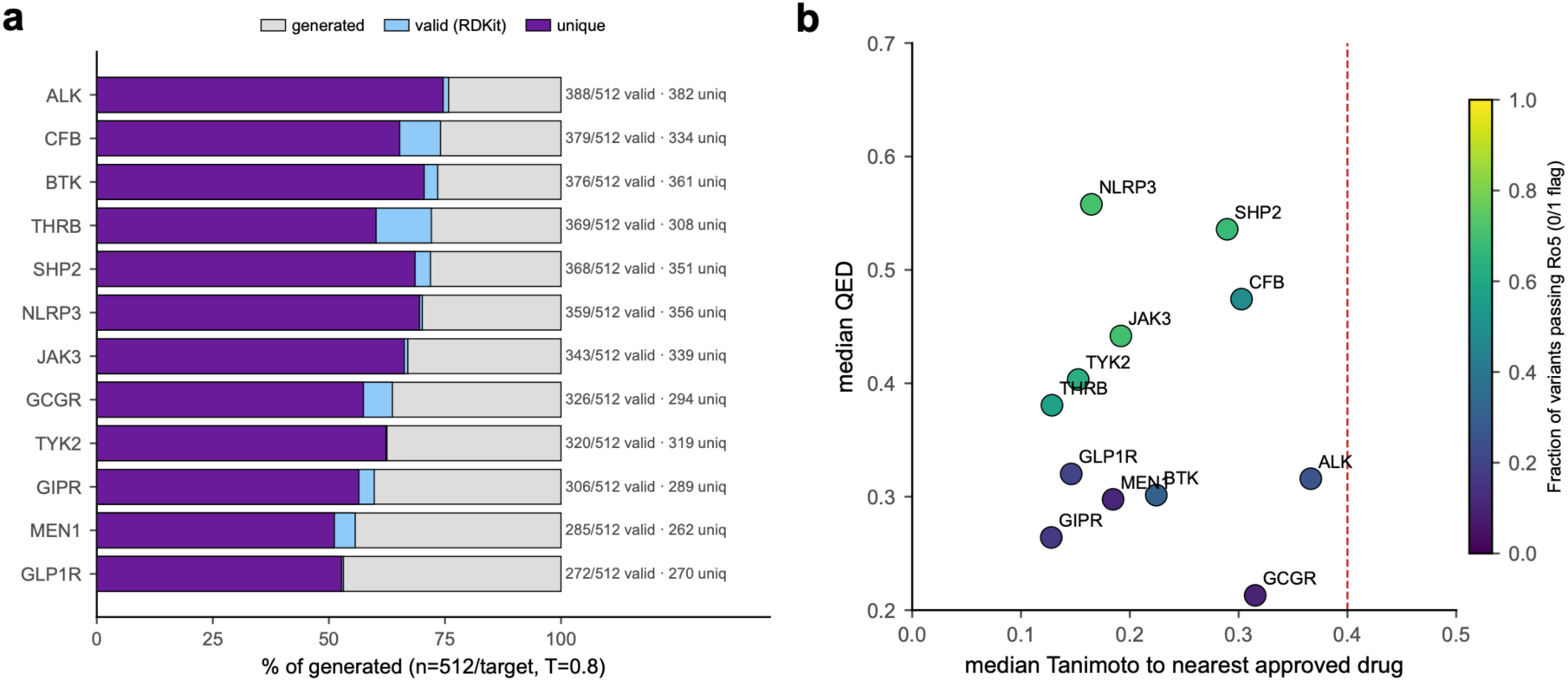
Decoder base-sampling quality across the 12 breadth targets. **(a)** Per-target generation throughput: of the n = 512 raw samples per target, the RDKit-valid and uniquely-canonicalised fractions. **(b)** Per-target starting chemistry: median ECFP4 Tanimoto to the nearest approved drug versus median QED for each target, coloured by the fraction of variants passing the Lipinski Rule-of-Five (0 or 1 flag); all targets fall below the 0.40 novelty threshold.

**Figure S2.**
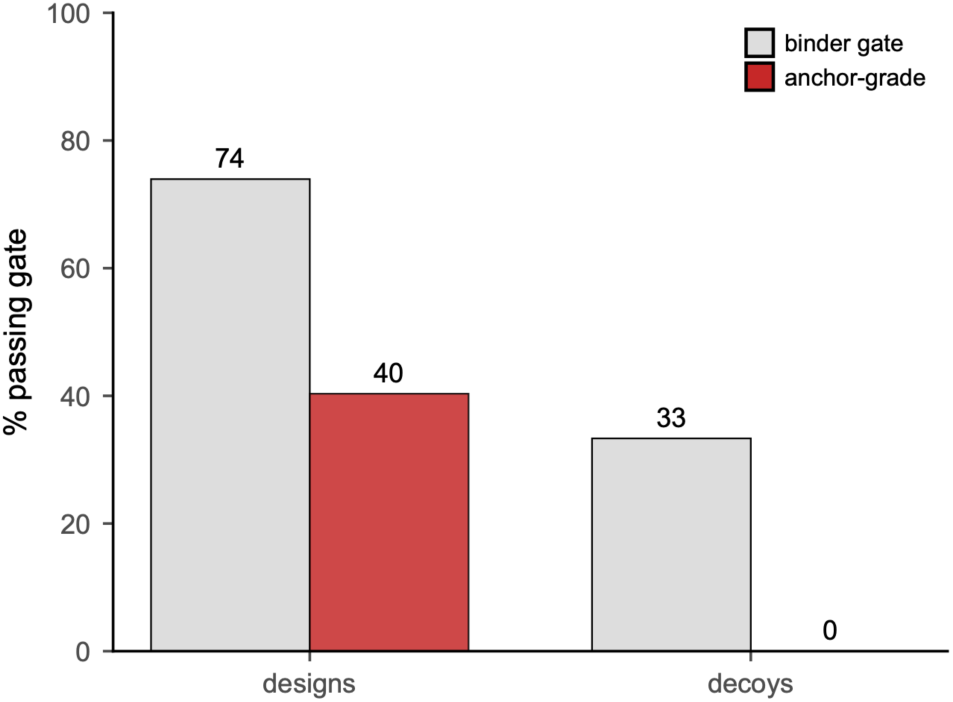
AF3 decoy-gate calibration. Negative-control decoy molecules cofolded against the 12 breadth targets; cohort-wise pass-rates at the MODERATE gate (iPTM ≥ 0.70, interface PAE ≤ 5 Å) and the STRONG gate (iPTM ≥ 0.90, interface PAE ≤ 1.67 Å) define the empirical false-positive floor.

**Figure S3.**
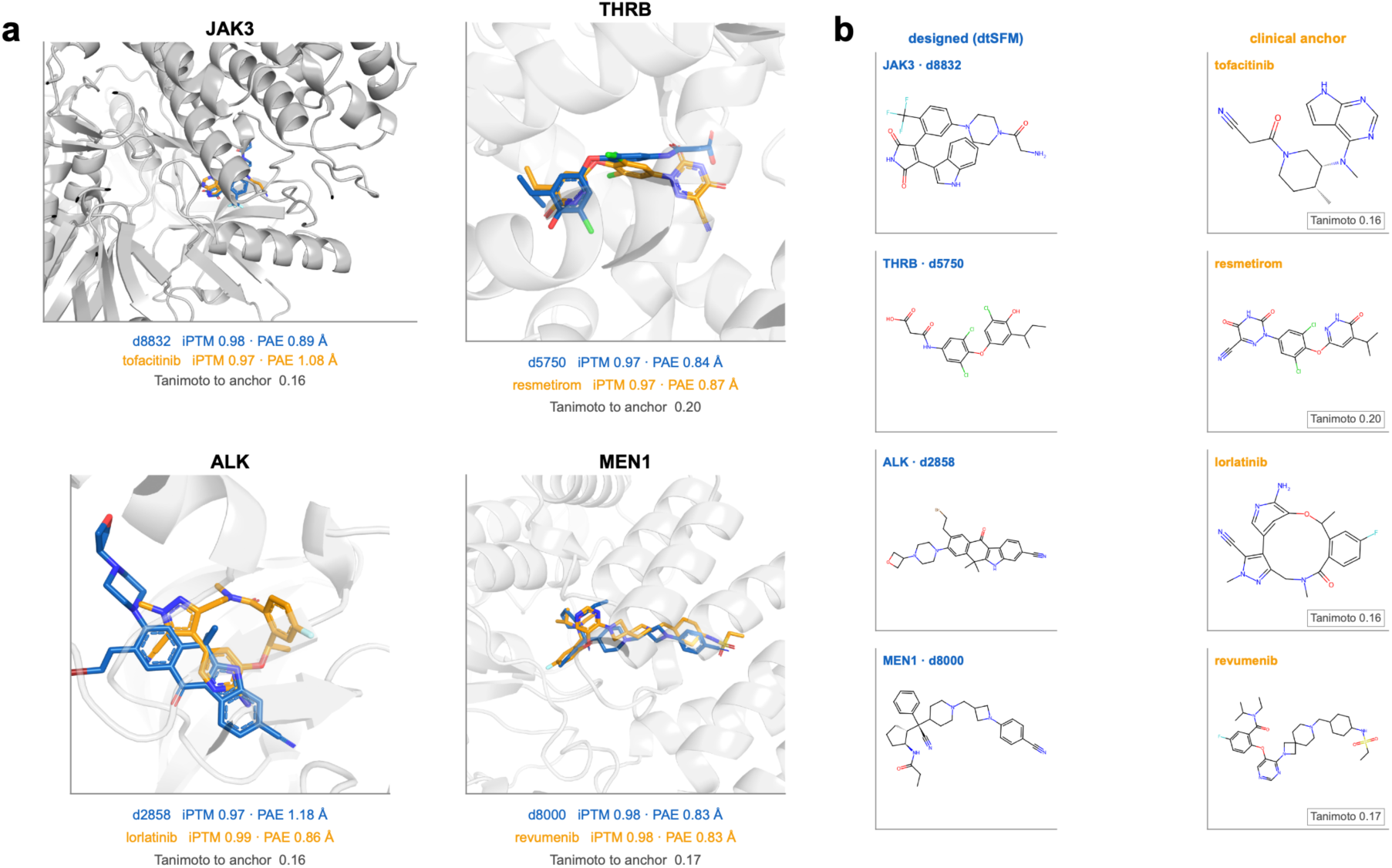
Structural and chemical characterization of the STRONG-tier breadth designs. **(a)** Representative four-cell gallery of AF3-predicted design–anchor pocket overlays for the clean STRONG-tier targets (JAK3, THRB, ALK, MEN1). **(b)** Two-dimensional chemistry pairs of the representative designs and their clinical anchors.

**Figure S4.**
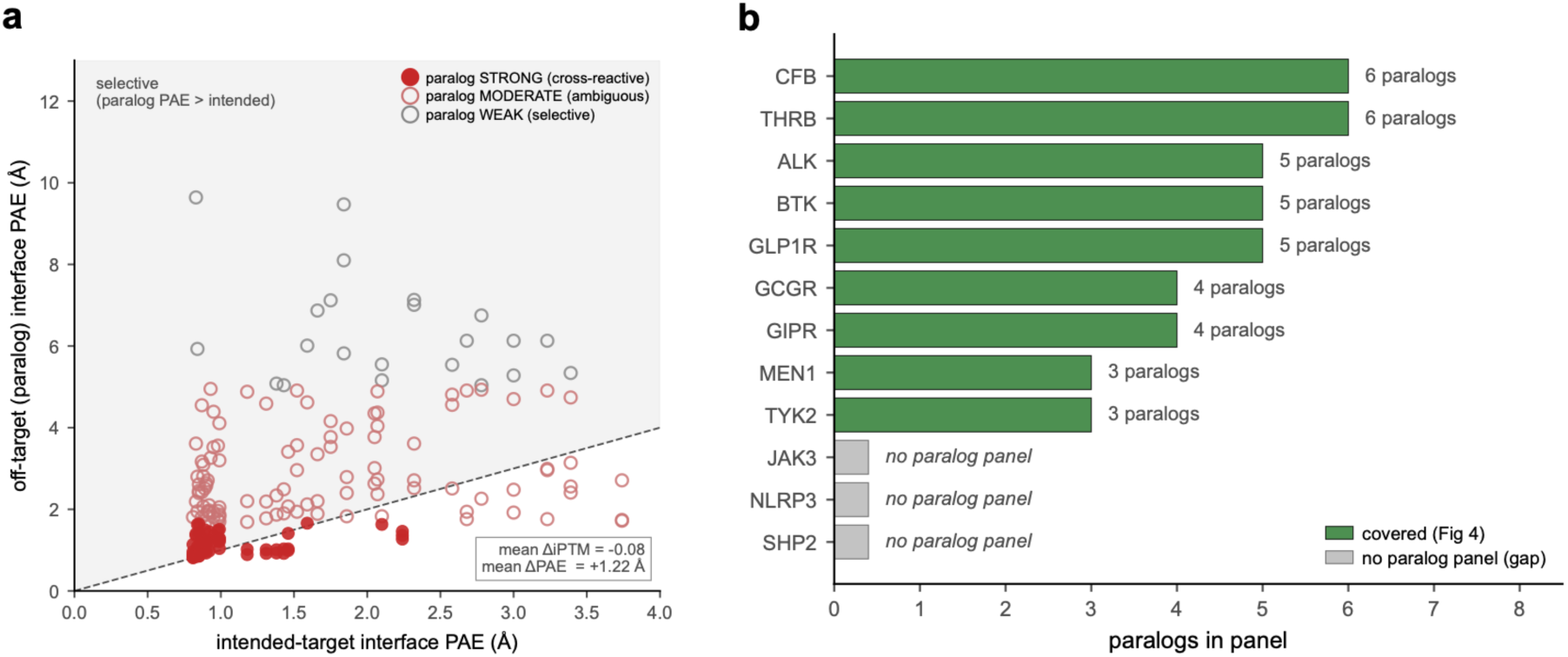
Paralog cross-reactivity profiling. **(a)** Paralog panel definition and extended cross-reactivity profiling across the breadth targets. **(b)** Paralog selectivity profiling for the breadth targets (JAK3, NLRP3, SHP were excluded).

**Figure S5.**
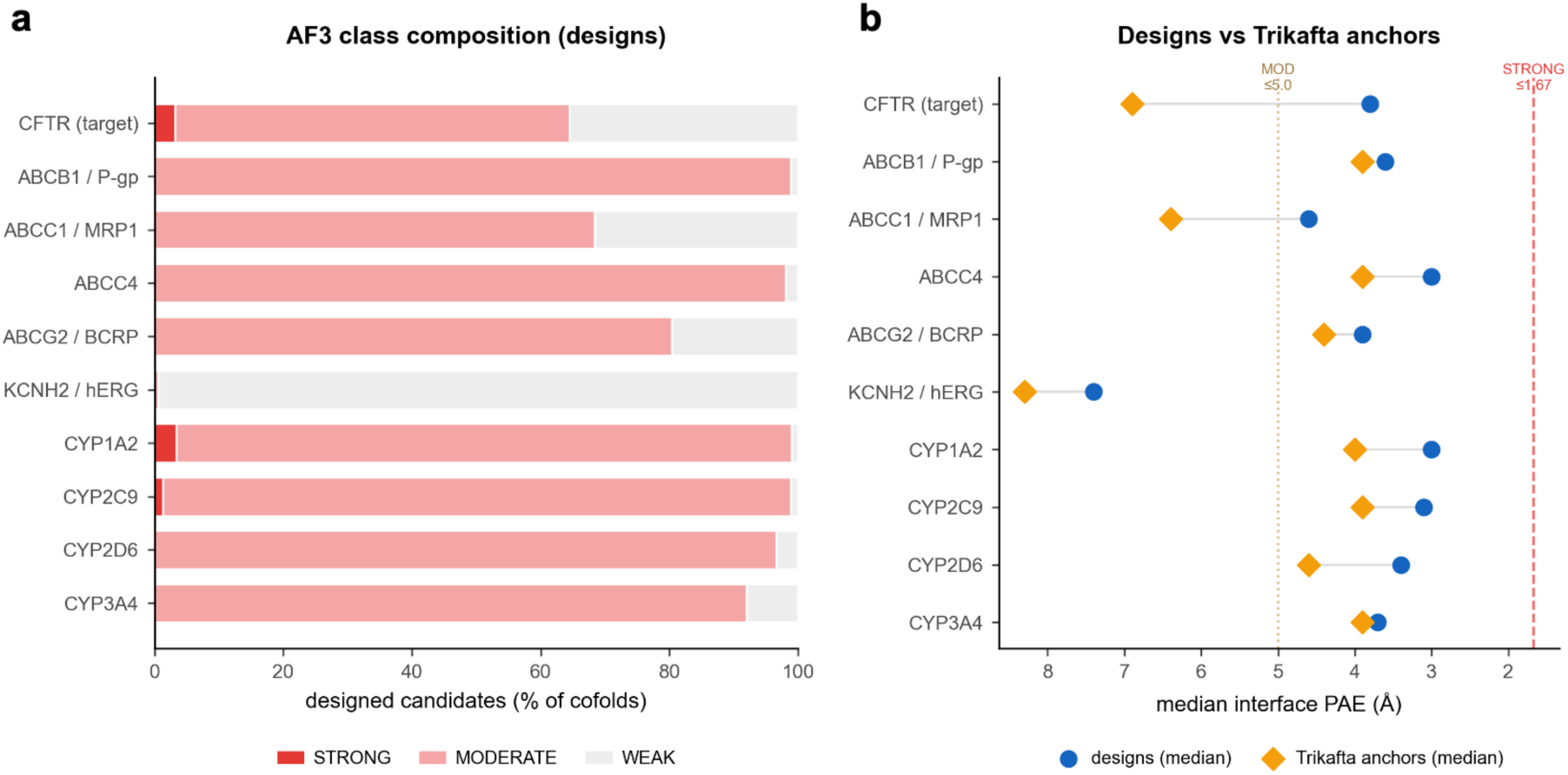
CFTR off-target panel. **(a)** AF3 class composition of designed candidates across the CFTR target, four ABC-transporter paralogs, and five safety-pharmacology proteins. **(b)** Designs versus Trikafta anchors: median interface PAE per protein at the decoy-calibrated thresholds.

**Figure S6.**
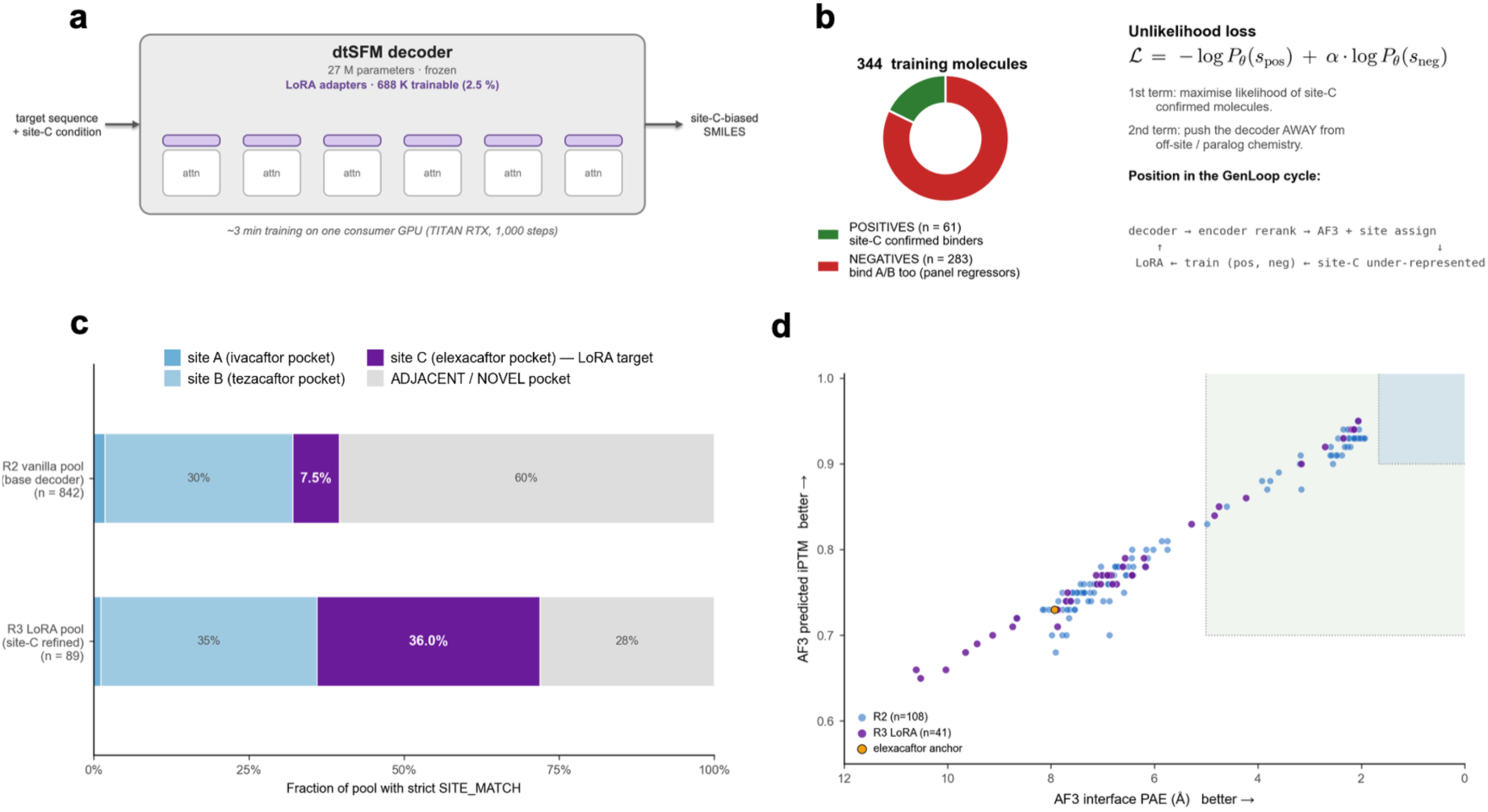
CFTR site-C LoRA directed-evolution refinement. (a, **b)** Site-C LoRA trained on 344 site-resolved molecules (61 site-C-verified positives versus 283 off-site negatives) under an unlikelihood loss that pushes the decoder toward the elexacaftor pocket and away from off-site negatives. **(c, d)** The refinement raised strict site-C SITE_MATCH from 7.5 % (n = 842) to 36.0 % (n = 89; 4.8-fold) while preserving AF3 confidence on the elexacaftor pocket (median iPTM 0.78 → 0.77, interface PAE 6.61 → 6.87 Å).

**Figure S7.**
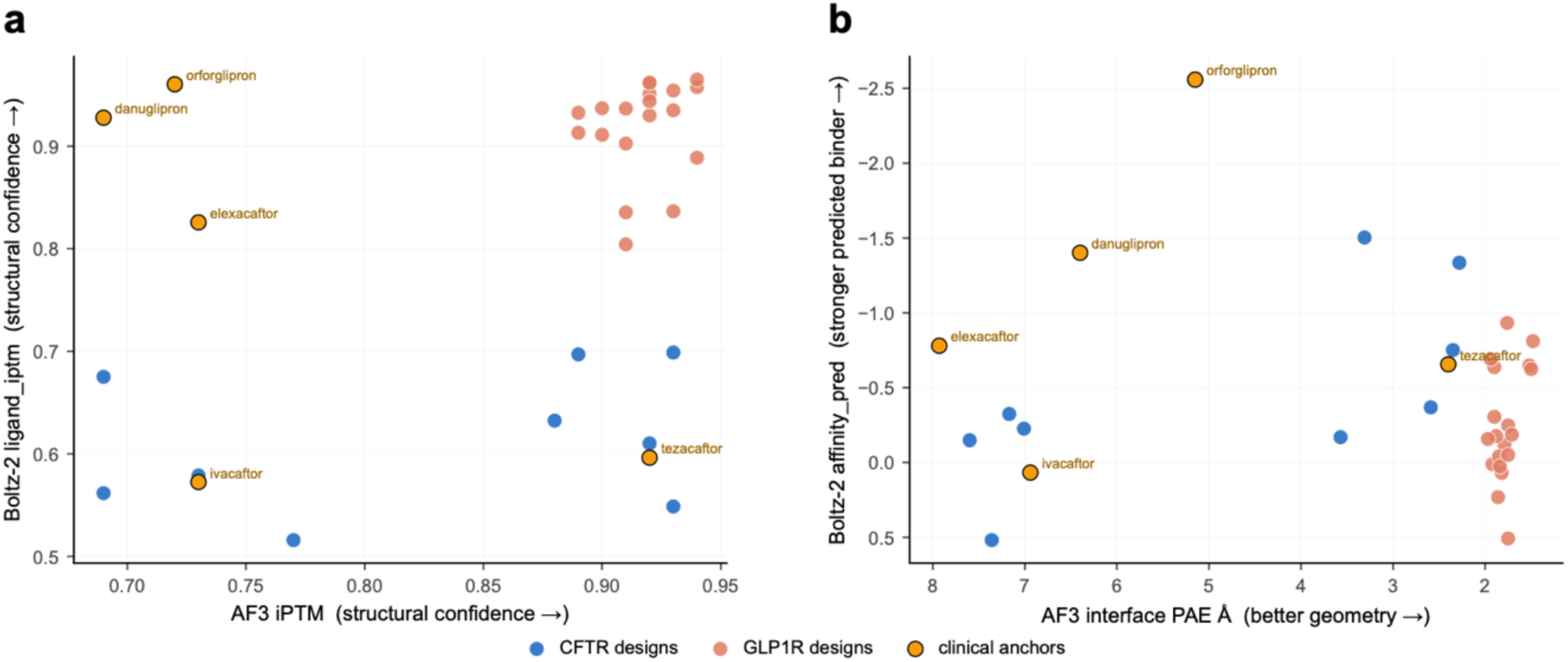
Cross-model structural verification (AF3 vs Boltz-2). Independent cofolding of the CFTR and GLP1R design cohorts and their clinical anchors by AF3 and Boltz-2. **(a)** AF3 iPTM versus Boltz-2 ligand_iptm (structural confidence); **(b)** AF3 interface PAE versus Boltz-2 affinity_pred (discriminating metrics, oriented so better is up and to the right). AF3 iPTM/PAE are structural-confidence metrics and Boltz-2 affinity_pred is a model-predicted (not measured) affinity; agreement reflects concordant structural modelling, not confirmed binding.

**Table S1.** Per-target AF3 outcome composition of designed candidates across the 12-target breadth screen. Tier: STRONG-tier ≥ 70 % STRONG, PARTIAL 5–30 %, ZERO 0 %.

| Target | Tier | n | STRONG | MODERATE | WEAK | %STRONG | Anchor |
| --- | --- | --- | --- | --- | --- | --- | --- |
| BTK | STRONG-tier | 18 | 18 | 0 | 0 | 100 | tirabrutinib |
| THRB | STRONG-tier | 21 | 20 | 1 | 0 | 95 | resmetirom |
| CFB | STRONG-tier | 21 | 20 | 1 | 0 | 95 | iptacopan |
| MEN1 | STRONG-tier | 20 | 15 | 3 | 2 | 75 | revumenib |
| ALK | STRONG-tier | 18 | 13 | 5 | 0 | 72 | lorlatinib |
| JAK3 | PARTIAL | 19 | 5 | 9 | 5 | 26 | tofacitinib |
| SHP2 | PARTIAL | 19 | 2 | 8 | 9 | 11 | SHP099 |
| GCGR | PARTIAL | 21 | 2 | 14 | 5 | 10 | LY2409021 |
| GIPR | PARTIAL | 21 | 1 | 16 | 4 | 5 | orforglipron |
| TYK2 | ZERO | 21 | 0 | 7 | 14 | 0 | deucravacitinib |
| NLRP3 | ZERO | 19 | 0 | 0 | 19 | 0 | MCC950 |
| GLP1R | ZERO | 20 | 0 | 16 | 4 | 0 | danuglipron |

**Table S2.**
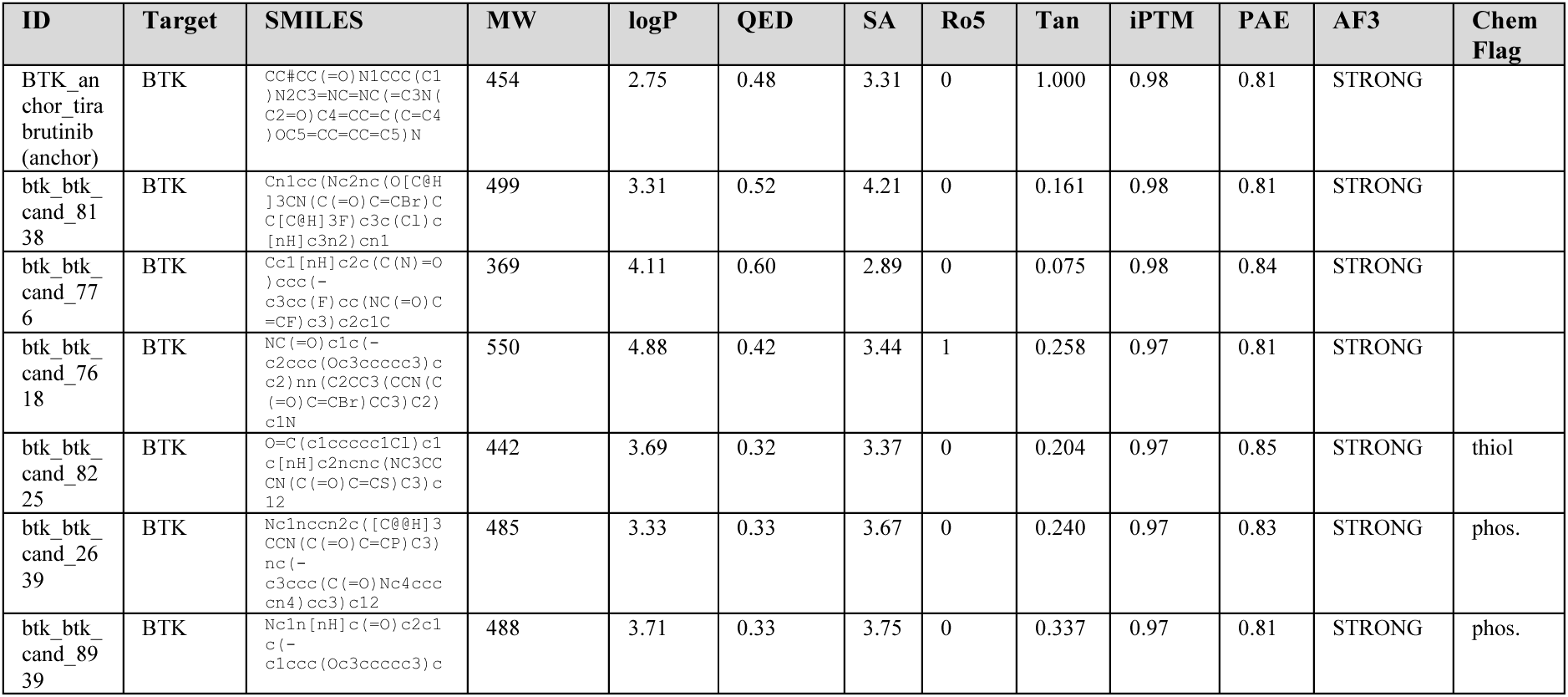
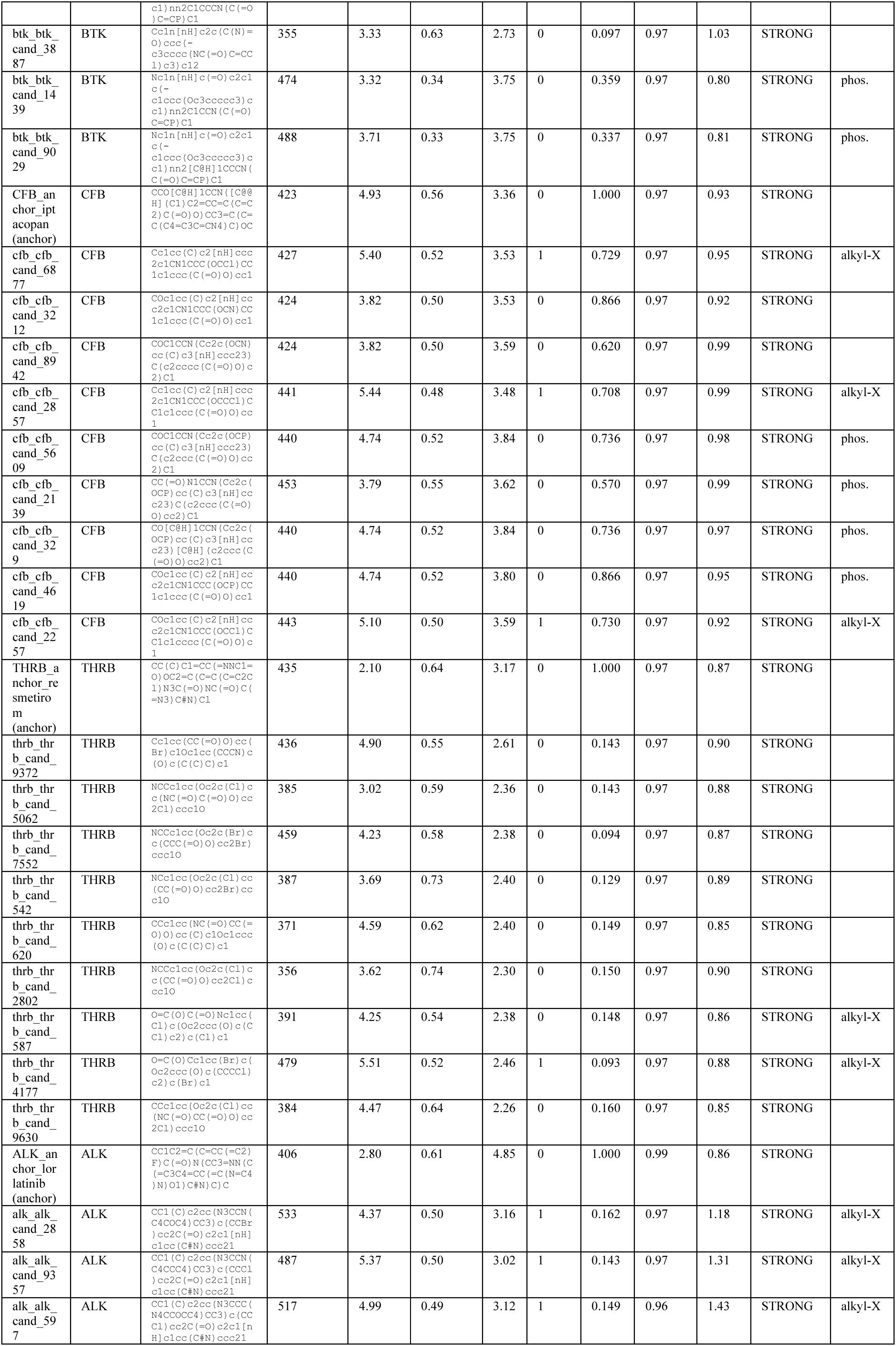
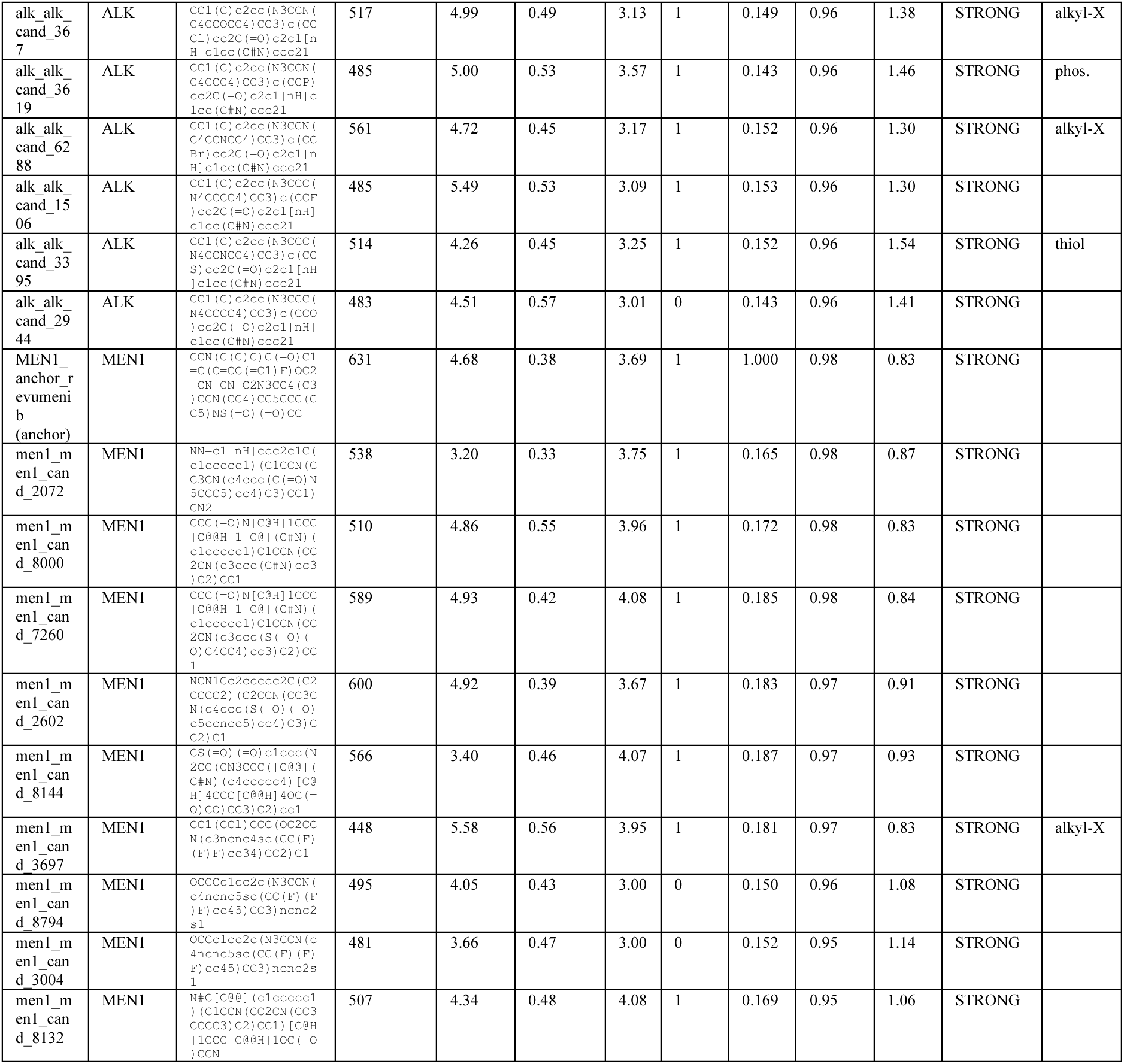
Per-candidate structural and chemical detail for the top-ranked designs (top 9 by AF3 iPTM) at the five STRONG-tier breadth targets, with each target’s clinical anchor. MW molecular weight; QED quantitative estimate of drug-likeness; SA synthetic accessibility; Ro5 Lipinski violations; Tan ECFP4 Tanimoto to the anchor; AF3 gate verdict; ChemFlag structural alert (alkyl-X / phos. / thiol; blank = none).

**Table S3.** Per-target paralog cross-reactivity composition and operational selectivity tier across the nine breadth targets with curated paralog panels (CLEAN < 25 % paralog STRONG, PARTIAL 25–75 %, FAMILY-SHARED > 75 %). Counts pool all candidate-versus-paralog AF3 cofolds per target.

| Target | Tier | n paralog cofolds | STRONG | MODERATE | WEAK | %paralog STRONG |
| --- | --- | --- | --- | --- | --- | --- |
| GCGR | CLEAN | 20 | 0 | 18 | 2 | 0 |
| GIPR | CLEAN | 20 | 2 | 11 | 7 | 10 |
| GLP1R | CLEAN | 25 | 0 | 19 | 6 | 0 |
| MEN1 | CLEAN | 15 | 0 | 10 | 5 | 0 |
| TYK2 | CLEAN | 15 | 3 | 9 | 3 | 20 |
| ALK | PARTIAL | 25 | 11 | 12 | 2 | 44 |
| CFB | PARTIAL | 30 | 14 | 16 | 0 | 47 |
| THRB | PARTIAL | 30 | 17 | 13 | 0 | 57 |
| BTK | FAMILY-SHARED | 25 | 24 | 1 | 0 | 96 |

**Table S4.**
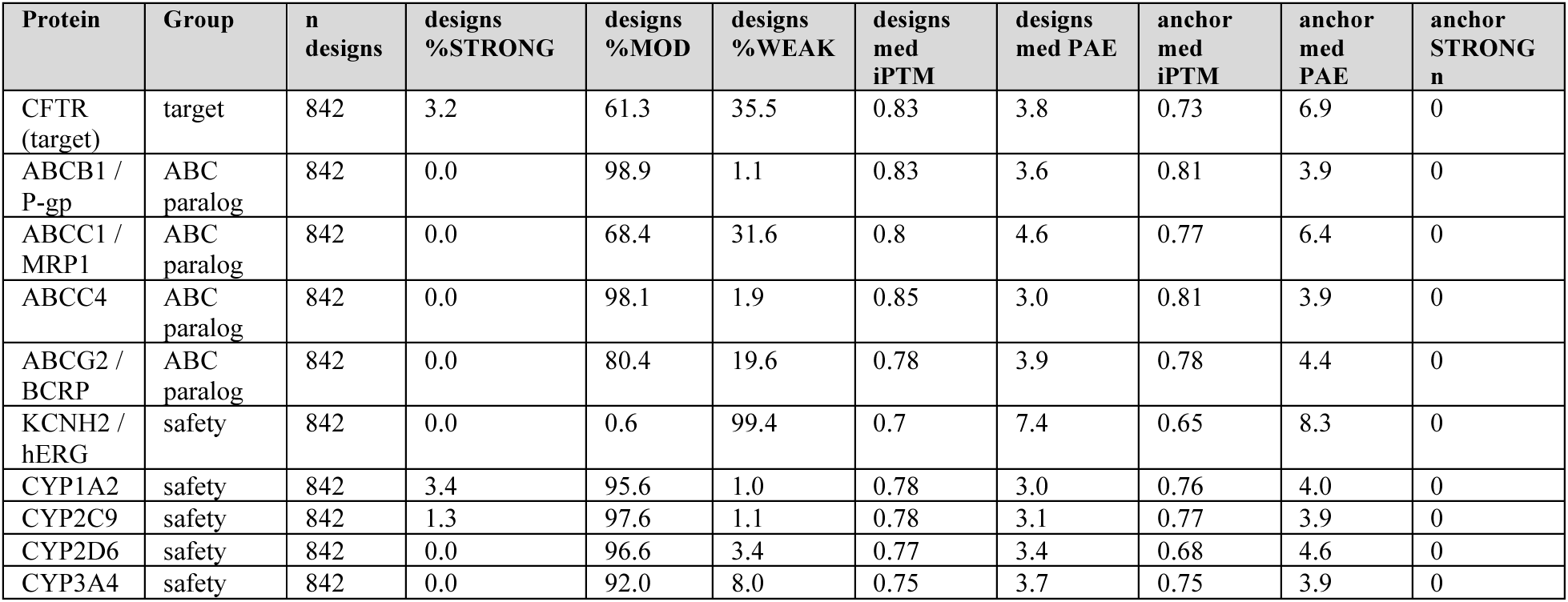
CFTR off-target panel: AF3 outcome composition of designed candidates versus the three Trikafta anchors across four ABC-transporter paralogs (ABCB1/ABCC1/ABCC4/ABCG2) and five safety-pharmacology proteins (hERG and four cytochrome-P450s), classified at the decoy-calibrated thresholds.

| Protein | Group | n designs | designs %STRONG | designs %MOD | designs %WEAK | designs med iPTM | designs med PAE | anchor med iPTM | anchor med PAE | anchor STRONG n |
| --- | --- | --- | --- | --- | --- | --- | --- | --- | --- | --- |
| CFTR (target) | target | 842 | 3.2 | 61.3 | 35.5 | 0.83 | 3.8 | 0.73 | 6.9 | 0 |
| ABCB1 / P-gp | ABC paralog | 842 | 0.0 | 98.9 | 1.1 | 0.83 | 3.6 | 0.81 | 3.9 | 0 |
| ABCC1 / MRP1 | ABC paralog | 842 | 0.0 | 68.4 | 31.6 | 0.8 | 4.6 | 0.77 | 6.4 | 0 |
| ABCC4 | ABC paralog | 842 | 0.0 | 98.1 | 1.9 | 0.85 | 3.0 | 0.81 | 3.9 | 0 |
| ABCG2 / BCRP | ABC paralog | 842 | 0.0 | 80.4 | 19.6 | 0.78 | 3.9 | 0.78 | 4.4 | 0 |
| KCNH2 / hERG | safety | 842 | 0.0 | 0.6 | 99.4 | 0.7 | 7.4 | 0.65 | 8.3 | 0 |
| CYP1A2 | safety | 842 | 3.4 | 95.6 | 1.0 | 0.78 | 3.0 | 0.76 | 4.0 | 0 |
| CYP2C9 | safety | 842 | 1.3 | 97.6 | 1.1 | 0.78 | 3.1 | 0.77 | 3.9 | 0 |
| CYP2D6 | safety | 842 | 0.0 | 96.6 | 3.4 | 0.77 | 3.4 | 0.68 | 4.6 | 0 |
| CYP3A4 | safety | 842 | 0.0 | 92.0 | 8.0 | 0.75 | 3.7 | 0.75 | 3.9 | 0 |

